# Population divergence manifested by genomic rearrangements in a keystone Arctic species with high gene flow

**DOI:** 10.1101/2024.06.28.597535

**Authors:** Siv N.K Hoff, Marius F. Maurstad, Alan Le Moan, Mark Ravinet, Christophe Pampoulie, Ireen Vieweg, France Collard, Denis Moiseev, Ian R. Bradbury, Ole K. Tørresen, Jane Aanestad Godiksen, Haakon Hop, Paul E. Renaud, Jasmine Nahrgang, Kjetill S. Jakobsen, Kim Præbel, Joël M. Durant, Sissel Jentoft

## Abstract

Genomic rearrangements have in recent years gained attention due to their evolutionary role in processes related to adaptation to local environmental conditions as well as diversification and speciation. In this study, we report on genomic rearrangements in the cold-water adapted polar cod (*Boreogadus saida*), a keystone Arctic fish species. By taking advantage of a new chromosome-level genome assembly in combination with whole-genome population sequencing data from specimens across the northern Barents Sea and adjacent regions, we identified a substantial number of larger chromosomal inversions (n=20) and characterized the previously identified chromosomal fusions (n=5). These genomic features — encompassing over 20% of the genome — exhibited genetic divergence, strong internal linkage disequilibrium, and signals of selection. Two of the identified inversions were associated with the two previously described hemoglobin clusters, while a third chromosomal region was found to differentiate between males and females. Moreover, clustering analyses on genotype frequencies of inversions revealed sub- structuring according to five geographic sub-groups suggesting sub-populations and/or the existence of cryptic ecotypes. These results provide novel insights into the impact of genomic rearrangements in population divergence and thus, potentially local adaptation, especially in species with high gene flow.

## Introduction

The genomic era has led to a growing body of literature demonstrating the importance of genomic rearrangements – such as chromosomal inversions and fusions – and their central role in evolutionary processes including diversification and adaptation to local environmental conditions^1–4^. Chromosomal inversions are mutations where a segment of the chromosome is reversed end to end, often containing multiple genes and regulatory elements. These large reorientations — if maintained within a population as variants — lead to reduced recombination between the two haplotype arrangements in the heterozygous state, resulting in higher linkage disequilibrium (LD) within the inverted region^5–7^. Such highly linked regions with different sets of inversion alleles allow for independent evolutionary trajectories of the two arrangements, making them important polymorphisms for species and populations and their ability to adapt to local environments despite gene flow. There are numerous examples across taxa where frequencies of polymorphic chromosomal inversions have been shown to vary in space and time, and thus, likely play essential roles in ecological adaptations to local environmental conditions^1–3,5,8,9^. Moreover, differences in life-history strategies, including timing of spawning and migratory behavior, are also reportedly linked to chromosomal inversions^10–14^. For instance, in Atlantic cod (*Gadus morhua*) the non-migratory Norwegian coastal cod (NCC) and the migratory Northeast Arctic cod (NEAC) are discriminated by three large chromosomal inversions, whereas the rest of the genome displays little genetic differentiation between the two ecotypes, indicating high degree of gene flow^10,11,15^. These findings further highlight the important role of chromosomal inversions, i.e., facilitating ecotype divergence, especially in species with high gene flow, which is the case for marine systems in general^12^.

Chromosomal fusions are another type of large-scale genomic rearrangement (similar to chromosomal inversions) that can change the recombination landscape across the chromosome, resulting in lower independence among loci within the given region^16^. As a result of reduced recombination within the fused region, it is thought that fusions may link advantageous loci together, similar to what is seen for chromosomal inversions^17,18^. Examples of identified chromosomal fusions that may play adaptive roles include the three-spined stickleback (*Gasterosteus aculeatus*)^19^ as well as Atlantic salmon (*Salmo salar*)^17^. Nonetheless, in marine systems, our understanding of the direct effects of chromosomal fusions on local adaptation and species complexes remains rudimentary.

The polar cod (*Boreogadus saida*) is a keystone cold-water adapted pelagic fish distributed across the Arctic Ocean and its shelf seas^20^. Despite its central role in the ecosystem, linking higher and lower trophic levels^21–23^, in-depth knowledge of its biology, migratory behavior, population structuring, and genomic composition in general is limited. For the Barents Sea area, previous studies have suggested that there are likely two spawning grounds, one in the Northwest of the Barents Sea and one in the southeast Barents Sea, adjacent to Novaya Zemlya^23–25^, Russia, suggesting that there could be at least two or more sub-populations within this region. Several studies have been carried out to characterize the population structure of polar cod throughout their circumpolar occurrence, most of which have utilized only a handful to hundreds of genetic markers, and the results have been inconsistent^26–29^. Limited population structure has been observed within the Barents Sea area^26–29^, however genetic differentiation between fjord populations on Svalbard and offshore Greenland specimens has been identified^27^. These results are indicative of possible local fjord populations, which is further supported by the evidence of local spawning populations and the differentiation of phenotypes between the western fjords of Svalbard^30–33^. When using hundreds of markers, no differentiation among polar cod sampled in the Barents Sea area was detected^29^. Notably, a handful of linked single nucleotide polymorphisms (SNPs) demonstrated genetic clustering, with a potential link to differences in size at specific age classes^29^.

To fully evaluate the population genetic structure and diversity of polar cod in the Barents Sea and adjacent regions we here take advantage of a newly generated chromosome-level genome assembly for polar cod (Hoff et al. submitted), in combination with a larger population level sequencing dataset (∼20X coverage) (Figure 1A). At the genome-wide level, we uncover limited population structuring, which indicates high levels of connectivity and gene flow in this region. Moreover, we detected numerous chromosomal inversions and fusions, displaying high degree of genomic differentiation, suggesting that these genomic rearrangements play an important role in local adaptation and/or ecotype divergence.

**Figure 1:**
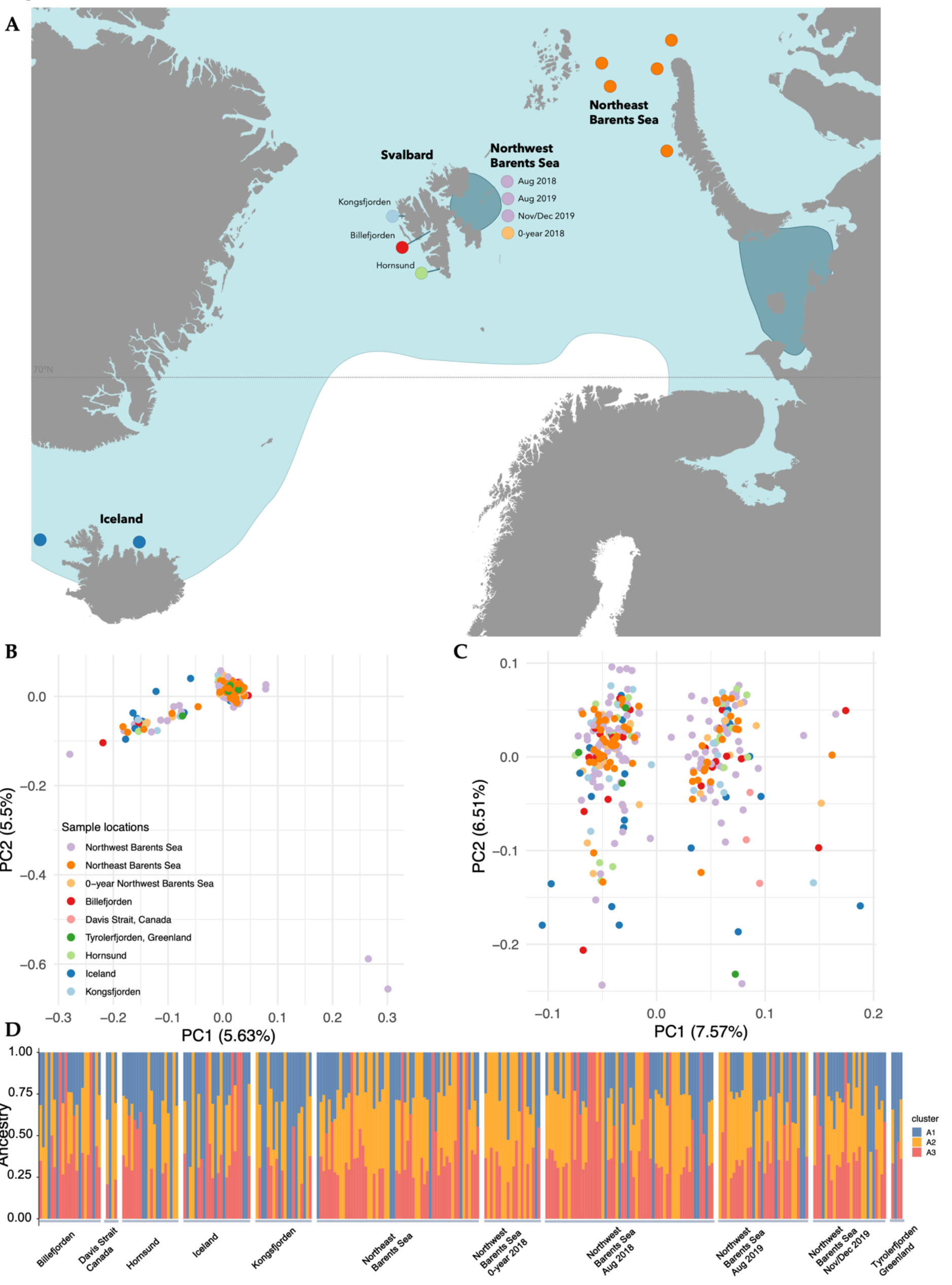
Whole genome population genetic structure of polar cod in the Barents Sea and adjacent regions and overview of sample locations. **A)** Map showing the distribution of polar cod in the northern Barents Sea region visualized in light blue, and the putative spawning regions marked in dark blue (redrawn from https://www.hi.no/en/hi/temasider/species/polar-cod). The sample locations of specimen collections used in the present study are marked as dots. Colors correspond to the legend in B). It should be noted that a few of the locations from where we have samples are not shown in this figure: Davis Strait, Canada, and Tyrolerfjorden, Greenland. PCAs across **B)** a reduced number of SNPs pruned for linkage and where high LD regions have been removed (n=564.970) and **C)** a set of high-quality SNPs (n=2.705.025) D) ADMIXTURE analyses (K=3) shown across the pruned genome-wide SNP set for the entire dataset (samples n=290).

## Results

### Genome-wide population structure and genetic diversity

The putative neutral SNP dataset (pruned for linkage as well as excluding all the highly linked regions), revealed a high level of intermixing and no strong signal of population structuring both for the PCA and ADMIXTURE^34^ analyses (Figure 1B, D, and S1). In the unpruned dataset, however, a clustering pattern along PC1 was observed and linked to the chromosomal rearrangements identified within the polar cod genome (see Figure 1C and S2 and section **Genome architecture and chromosomal inversions** below).

The genomic differentiation estimated as pairwise fixation index (FST)^35^ between samples collected in fjords of Svalbard, Northwest, and Northeast Barents ranged between ∼ 0 and 0.00195 (see Table S1 for an overview of global FST estimates between locations, and Supplementary html: FST), further supporting the overall low differentiation across the locations included in this study. For instance, we did not detect greater differences between polar cod caught at more distant locations, such as between Northeast and Northwest Barents Sea, than between specimens caught within one geographic location (such as the Northwest Barents Sea) at different seasons and/or years (Table S1). High genome-wide differentiation was detected when samples from the Barents Sea or the fjords were compared to Davis Strait, Canada (Table S1), with the highest FST estimate of 0.016 between Northwest Barents Sea summer 2018 and Davis Strait (Table S1). These results corroborate previous studies conducted on smaller sets of nuclear as well as mitochondrial markers, showing low levels of spatial population structure among polar cod sampled at a relatively large spatial scale^28,36^ as well as in the Barents Sea and Arctic Ocean region^26,29^. Based on two hypothesized spawning grounds in the Barents Sea, i.e., in the northwestern Barents Sea and southeastern Barents Sea area^23–25^, our findings suggest that natal homing behavior is unlikely to be prominent, and rather indicates that there is high connectivity between different locations and/or sub-populations within the Barents Sea area.

### Genome architecture and chromosomal inversions

#### Identification of 20 chromosomal inversions and their characteristics

Inspections of local PCAs in combination with pairwise *r*^2^ estimates (i.e., indicating patterns of LD, see Figure S3), as well as evaluation of heterozygosity levels along the chromosomes, enabled the identification of a total of 20 putative chromosomal inversions across the polar cod genome. These regions were found to range from ∼3 Kb to ∼15 Mb in size (spanning at least 15% of the genome of polar cod, see Figure 2A and Table S2) and contain 29 to 974 genes. Similar approaches have previously been applied and used to uncover inversions in various taxa^1,3,9,37–40^. Of the inversions uncovered in polar cod, 11 displayed a classical inversion pattern with three distinct clusters representing the homokaryotypes (of each arrangement), the heterokaryotypes (see Figure 3 and Supplementary html: classical inversions), for further description of characteristics and identification of inversions see^1,3,9,37–40^. Additionally, while conducting PCAs of SNPs within inversion regions we identified a second type of inversion, which displayed a more complex clustering pattern for one of the two homozygous arrangements. For some of these inversions, one of the homokaryotypes as well as heterokaryotypes displayed a dichotomous or continuous pattern of variation along PC2 (examples include Bschr7.01and Bschr12.02; see Figure S4). Moreover, for some of the most extreme cases of these complex inversions, additional subclusters were observed resulting in three distinct homokaryotype clusters, and where heterokaryotypes between all of the homokaryotypes were detected (exemplified by Bschr16.02; Figure 4 and Figures S5 and S6). All inversions detected, their placement on the respective chromosomes, and number of genes within them are listed in Table S2. For more descriptions of the inversions, see Supplementary Note 1.

**Figure 2:**
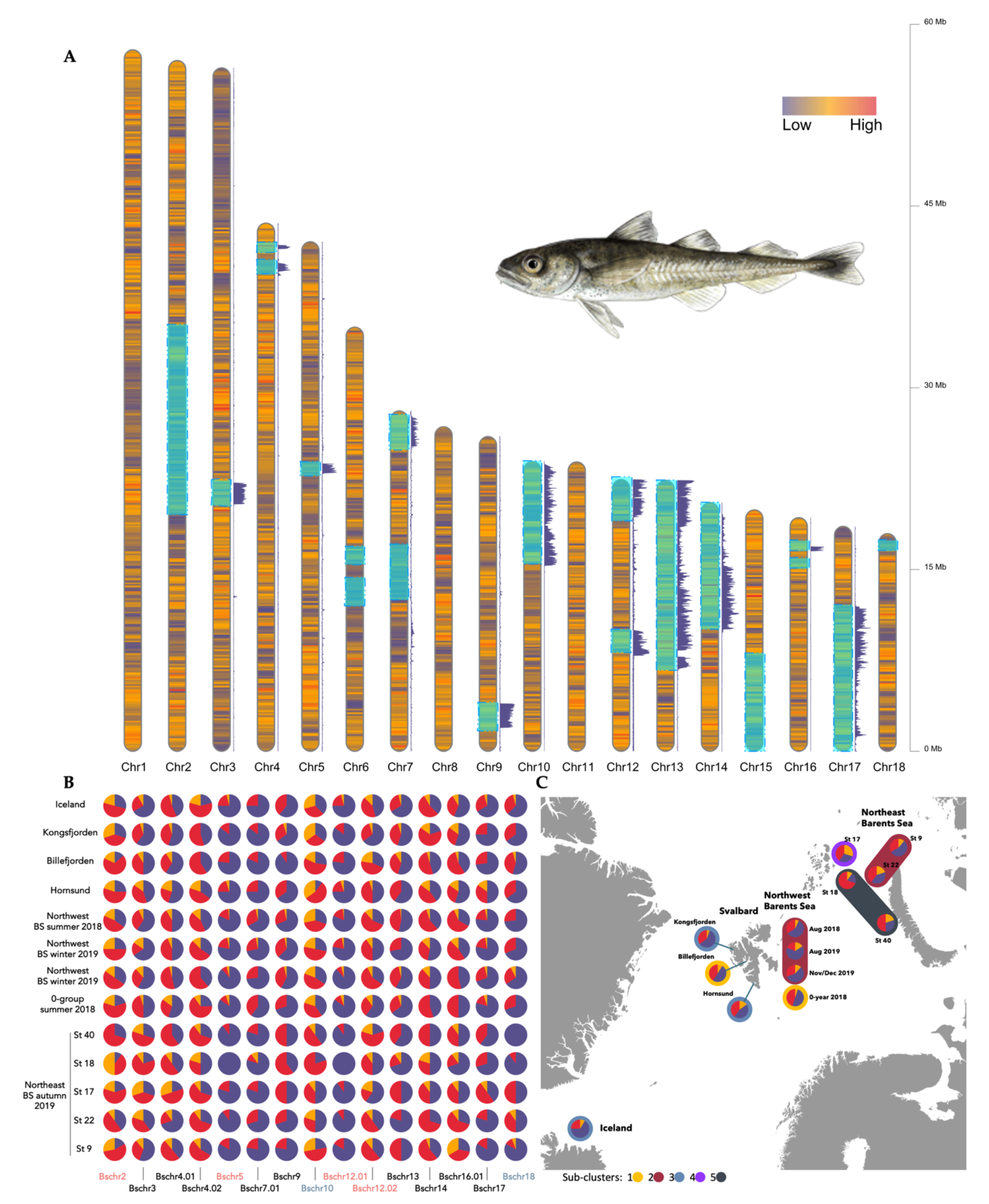
The polar cod genome is characterized by many chromosomal inversions. **A)** Schematic overview of chromosomal inversions detected in the present study, with pairwise FST estimates between the two homokaryotypes visualized along the chromosomes. The density map displays the frequency of coding sequence (CDS) in 150000 bp windows along chromosomes**. B)** Genotype frequency of 15 of the 20 inversions across locations and seasons in the northern Barents Sea region. Genotypes shown as Yellow=Hom rare, red=Het and purple=Hom common. Colored names (red or blue) indicate inversions found to be in linkage **C)** Genotype frequency distribution of the classical inversion Bschr3 and sample locations colored according to the five delineated sub-clusters using genotypes of 19 inversions. Illustration of polar cod (*Boreogadus saida*) by Alexandra Viertler.

**Figure 3:**
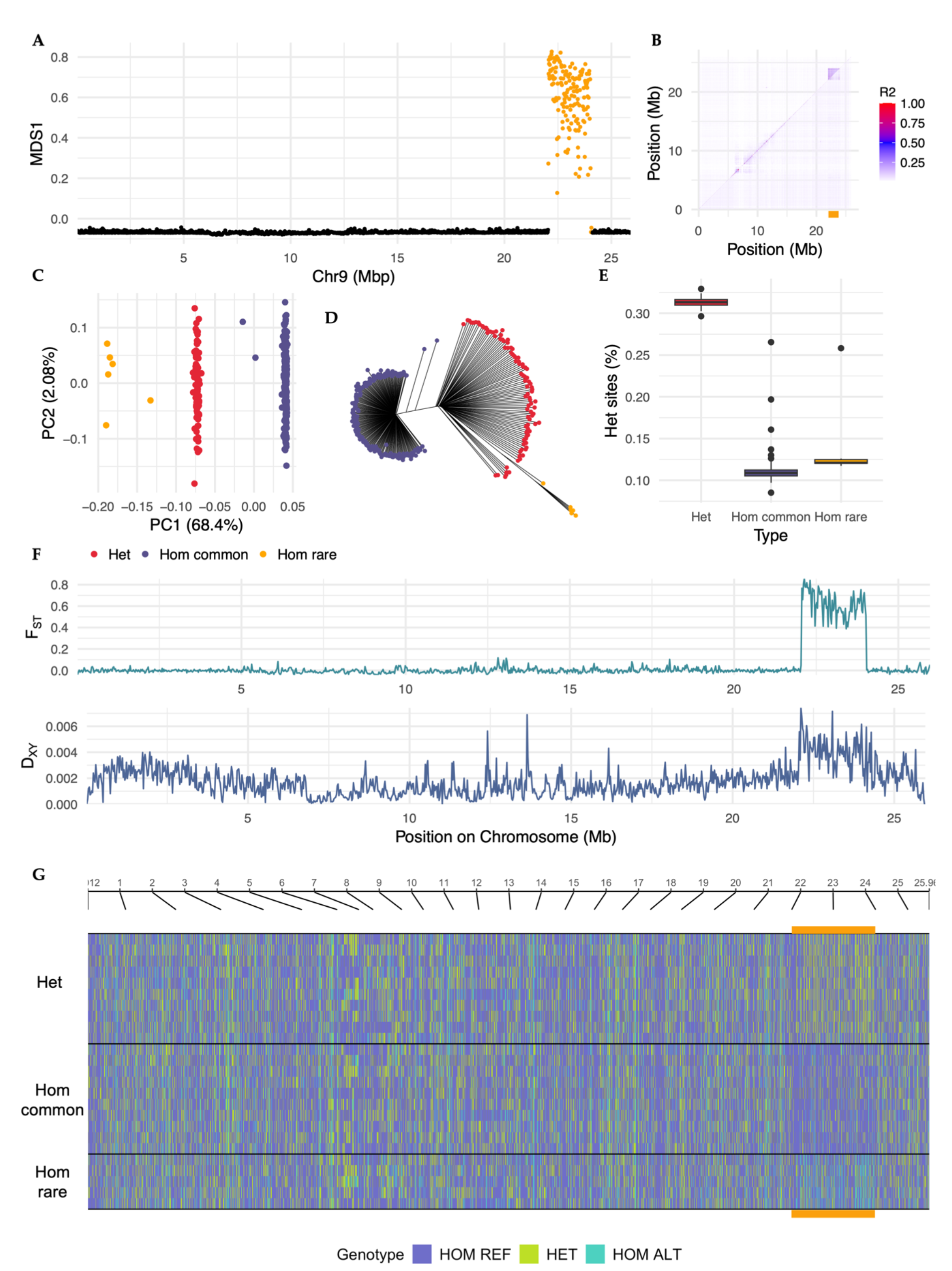
Characteristics of a classical inversion: Bschr9. **A)** MDS1 variation along chromosome 9 generated by lostruct^91^. **B)** *r*^2^ heatmap along chromosome 9, upper triangle calculations were done among all 290 specimens, whereas lower panel only among Hom common samples. The location of Bschr9 is marked with a yellow bar. **C)** PCA of SNPs residing within the inversion region. **D)** Neighbor-joining clustering of SNPs residing within the inversion region. **E)** PCA of SNPs residing within the inversion region with statistics for heterozygous sites (%) visualized as a color scale. **F)** Pairwise FST and DXY (in 25Kb windows along chromosome 9) comparisons between individuals representing each homozygous cluster. **G)** SNP genotype composition along the chromosome, of a selection of samples of each genotype determined (i.e., being classified as either HOM ALT, HOM REF, or heterozygous (HET) in accordance with the reference genome), with the inversion location marked in yellow.

**Figure 4:**
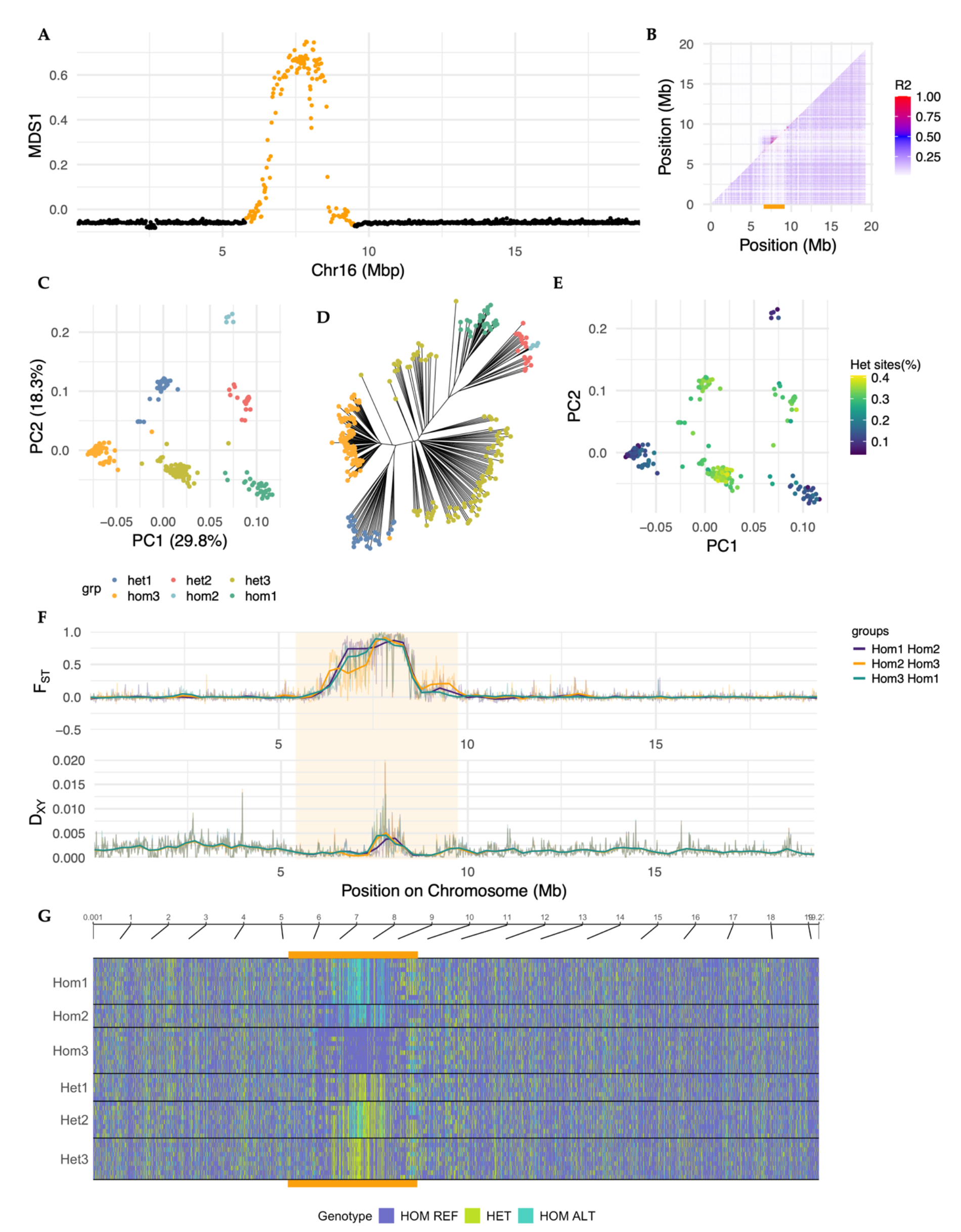
Characteristics of a complex inversion: Bschr16.02. **A)** MDS1 variation along chromosome 16 generated by lostruct^91^. **B)** *r*^2^ heatmap along chromosome 16, upper triangle contains all 290 fish, whereas lower panel only among a selection of Hom3 samples. Location of Bschr16.02 is marked with a yellow bar. **C)** PCA of SNPs residing within the inversion region. **D)** Neighbor-joining clustering of SNPs residing within the inversion region. **E)** PCA of SNPs residing within the inversion region with statistics for Het sites (%) visualized as a color scale. **F)** Pairwise FST and DXY (in 25Kb windows along chromosome 16) comparisons between individuals representing each homozygous cluster. **G)** SNP genotype composition along the chromosome, of a selection of samples of each genotype determined (i.e., being classified as either HOM ALT, HOM REF, or heterozygous (HET) in accordance with the reference genome), with the inversion location marked in yellow.

The pairwise FST (estimated along chromosomes) between the homokaryotypes for the classical as well as for some of the complex inversion types (Bschr16.02, Bschr6.02, and Bschr7.02) revealed high genetic differentiation between the two or three homokaryotypes for classical and complex inversions, respectively (Figure 2A and Figure 3F for classical inversions, and Figure 4F, Figure S5D and Figure S6D for complex inversions). For some of the complex inversion types, e.g., Bschr16.02 and Bschr7.02, pairwise FST estimates uncovered differences in signals of divergence in the boundary regions between the three homokaryotypes (Figure 4F and Figure S6D), which could suggest length variation between haplotypes. Additionally, for Bschr7.02 differential patterns of divergence were detected at the center of the inversion between the three homokaryotypes, further confirmed by the genotype calling (Figure S6D and E).

The detailed inspection of SNP genotype variants called (i.e., classified as either homozygote for the reference allele (HOM REF), homozygote for the alternative allele (HOM ALT), or heterozygous (HET)) along the chromosome provided further insight into the genotype composition of the different inversion types. For instance, within the two homozygous haplotypes for the classical inversion Bschr9, we detected the region displaying mainly HOM REF or HOM ALT variants, while more HET variants were present within the heterozygous haplotype (Figure 3G), as expected. For three of the complex inversions that we inspected in more detail (i.e., Bschr6.02, Bschr7.02 and Bschr16.02), however, we found that the three homozygous haplotypes (Hom1, Hom2 and Hom3) were characterized by unique ’’blocks’’ of more HOM ALT, HOM REF, or heterozygous variants (Figure 4G, Figure S5D and Figure S6D). For Bschr16.02, we found that Hom3 was primarily composed of HOM REF genotypes. Moreover, Hom1 harbored mainly the HOM ALT genotypes in the beginning, and more HOM REF at the end of the region. Hom2 constituted of primarily HOM REF genotype at the beginning of the inversion region, more HOM ALT genotypes in the middle of the region, and primarily HOM REF at the end of the region (Figure 4G). Furthermore, at the very end of this region, Hom1 and Hom3 displayed more HET genotypes, while Hom2 seemed to harbor more of the HOM REF genotypes (Figure 4G). These findings combined, suggest either length differentiation and/or the potential of one or several nested inversions inside the defined inversion region, which are in line with complex inversion structures also described in Australian zebra finch (*Taeniopygia guttata castanotis*)^41^ as well as rough periwinkle (*Littorina saxatilis*)^42^.

#### Frequency and distribution of inversion genotypes across spatial and temporal scales

The distribution of the inversion genotypes (i.e., of the 15 identified classical inversions, Table S2) uncovered differences in frequencies of the inversion genotypes across sampling locations as well as between seasons (Figure 2B). For Bschr5, Bschr7.01, Bschr9, Bschr12.01, Bschr13, Bschr17, and Bschr18, an overrepresentation of one homokaryotype (Hom common, see Materials and Methods for naming of the genotypes) was observed within the study area, while only a few samples were found to be of the alternate homozygous genotype (Hom rare) (Figure 2B). Conversely, for Bschr2, Bschr3, Bschr4.01, Bschr4.02, Bschr10, Bschr12.02, Bschr14 and Bschr16.01 the frequencies of the three inversion karyotypes (Hom common, Het and Hom rare) were more balanced and/or excess of heterokaryotypes across most of the sampling sites (Figure 2B).

Intriguingly, we also observed fine-scale differences in the frequencies of several of the inversions on a spatial as well as temporal scale (Figure 2B and C). For instance, the Bschr9 Hom common variant (Figure 2B) was found in higher frequency in Billefjorden compared to the other sampling locations (Figure 2B), and the inversion is detectable in pairwise FST (measured along chromosomes) comparisons between Billefjorden and any other location (see Supplementary html: FST). For Bschr3, Hom rare and heterozygous individuals were observed in higher frequency in Northeast Barents Sea sample locations (St 40, 18, 17, and 22) compared to northwestern sampling locations (Figure 2B and C). For Bschr4.02 no Hom rare individuals were detected in Kongsfjorden, Billefjorden nor the Northwest Barents Sea winter 2019 samples (Figure 2B). For the highly linked inversions, Bschr5 and Bschr12.01 (see details below), no Hom rare individuals were found among Northeast Barents Sea samples as well as Kongsfjorden and Billefjorden (Figure 2B). For Bschr10 very few Hom common individuals were observed in Hornsund samples, while two of the Northeast Barents Sea locations have a higher frequency of Hom common (sample stations St 17 and 22, see Figure 2B). Additionally, for one of the inversions, Bschr14, frequency differences between age classes were detected (Figure S8). More specifically, for some sampling locations (i.e., Northwest Barents Sea (summer and autumn 2019), Kongsfjorden, and Billefjorden) Hom rare became less prominent in 2- year-old vs. 1-year-old individuals, whilst the other inversions retained a more balanced karyotype frequency throughout the year-classes (Figure S7 and Supplementary html: classic inversions).

When testing for correlation between the different inversions and their genotypes, we found that Bschr5 and Bschr12.01 were highly linked. Moreover, we detected linkage between Bschr5 and Bschr12.02 (5.7%), between Bschr12.01 and Bschr12.02 (5.6%), between Bschr18 and Bschr10 (3.95%), and lastly, between Bschr2.02 and Bschr12.02 (2.4%) (Figure 2B and Table S3). Due to the high degree of complexity for several of the inversions, we only performed the analyses for the inversions displaying three distinct karyotype clusters (n=15), thus additional linkage between some of the inversions cannot be ruled out. When treating the entire dataset (all 290 polar cod samples) as one population, we detected that some of the inversions showed deviation from Hardy Weinberg equilibrium^43^ (HWE) (p-value < 0.05). This included Bschr3, Bschr4.02, Bschr5, Bschr12.01 and Bschr14 (Table S4). For inversions Bschr3, Bschr5, and Bschr12.01, the deviation from HWE is likely attributed to heterozygous deficiency, with positive FIS estimated range from 0.128 to 0.138 (Table S4). Conversely, for Bschr4.02 and Bschr14 a heterozygous excess was observed with FIS estimations of -0.138 and -0.128, respectively (Table S4).

#### Identification of other high LD regions

In addition to the chromosomal inversions, we identified regions displaying elevated *r*^2^ and FST (estimated between sample locations) in the middle of the larger chromosomes, i.e., Chr1, Chr2, Chr4, and Chr5, compared to the distal parts of the chromosomes (Figure S3 and Supplementary html: FST). For most of these regions, we observed lower estimates of π (nucleotide diversity) as well as DXY (nucleotide divergence) within vs. outside these genomic regions (Supplementary html: FST). These regions were found to overlap with the fused regions and localization of the centromere within these chromosomes^44^. Additionally, we highlight that for the fused central region on chromosome 2, a complex inversion was detected (Bschr2), further indicating a potential evolutionary role of these fused regions, by linking adaptive genes together.

#### Signals of selection along chromosomes

By investigating signals of selection along the chromosomes, we detected positive estimates of Tajima’s D^45^, especially in genomic locations surrounding breakpoints of some of the chromosomal inversions identified (Figure S8 and Supplementary html: Tajima’s D). Most of the inversions displayed overall neutral or slightly positive estimates for Tajima’s D along its length (Figure S8). However, inversions Bschr6.01, Bschr7, Bschr9, Bschr12.01, and the end region of Bschr15, displayed negative estimates of Tajima’s D (Figure S8). Notably, for many of the identified inversions, local regions within the inversion either the ends (breakpoints), one of the ends, or the central region were found to display unique signals of selection. For instance, Bschr2 and Bschr16.02 exhibit valleys of negative estimates of Tajima’s D at the breakpoint regions, whereas the region between displays slight positive estimates (Figure S8 Supplementary html: Tajima’s D). Moreover, Bschr6, Bschr7.02, and Bschr12.02 are examples where higher estimates of Tajima’s D were detected in one breakpoint region compared to the other (Figure S8, Supplementary html: Tajima’s D). Interestingly, inversion Bschr2, which likely overlaps with a fusion region on chromosome 2 (see above section of regions displaying elevated LD for more details), displayed positive values of Tajima’s D along its length, but negative values in both boundary regions (Figure S8 and Supplementary html: Tajima’s D). Additional tests of signals of selection by integrated haplotype score^46^ (iHS) and cross-population extended haplotype homozygosity^47^ (XP-EHH) score show similar trends as described above (for more details see Supplementary note 2 and Supplementary htmls: iHS and XP-EHH), and to some extent, but less clear, for the calculated nucleotide diversity estimates (π) (for details see Figure S9 and Supplementary note 2).

#### Hemoglobin gene clusters and chromosomal inversions

A manual inspection of the hemoglobin gene clusters revealed the MN (i.e., *mpg* and *nprl3* genes) and LA (i.e., *lcmt1* and *aqp8a* genes)^48^ hemoglobin clusters to be located on chromosomes 12 and 15, respectively. The MN cluster, from the upstream flanking gene *nprl3* to the downstream flanking gene *kank2* was found to be located Chr12: 14722709 – 14744959 (Table S5), in close proximity (∼45.000 bp downstream) to inversion Bschr12.02 (Figure 6). Intriguingly, the LA cluster was found to be located within inversion Bschr15, at position 14153907 – 14206026 (Figure 6 and Table S5).

#### Detection of substructure among the samples using inversion genotype frequencies

By utilizing the inversions as genetic markers, we explored signals of sub-clusters among our samples. The results delineated our samples into five different groups based on similarity in genotype frequencies (Table 1 and visualized on Figure 2C). The Icelandic samples grouped with the fjords of Svalbard samples, except for fish caught in Billefjorden, which grouped with the Barents Sea 0-group samples from 2018. The Barents Sea samples were found to display similar genotype frequencies leading to one group of Northwest Barents Sea even though samples were collected across different years (i.e., summer 2018, summer 2019, and winter 2019). Moreover, the analyses further grouped the Northwest Barents Sea together with two Eastern Barents Sea sampling locations north of Novaya Zemlya/Kara Sea (Table 1). From the remaining three Eastern Barents Sea sample locations, two groups were identified, one group containing Eastern Barents Sea location St 17, and another group with Eastern Barents Sea location St 18 and St 40 (Table 1). Using these sub-clusters for testing the inversion genotypes frequencies did not increase the number of inversions deviating from HWE vs. testing all samples as one meta-population (Table S6), supporting that the sub-clusters identified are biologically meaningful.

**Table 1.** Clustering analyses based on inversion genotype frequency and distribution.

| Cluster | Sample locations included in the sub-clusters | Chromosomal inversion(s) decisive for the sub-clusters |
| --- | --- | --- |
| 1 | 0-year Northwest Barents Sea + Billefjorden | Bschr3, Bschr6.02, Bschr16.02 |
| 2 | Northwest Barents Sea (summer 2018, summer 2019 and winter 2019) + Northeast Barents Sea (station 22 + 9) | Bschr6.01, Bschr16.01 |
| 3 | Hornsund + Iceland + Kongsfjorden | Bschr18 |
| 4 | Northeast Barents Sea (station 17) | Bschr18, Bschr12.01, on Bschr16.01, Bschr17, Bschr3, Bschr12.01, Bschr7.02, Bschr16.02, Bschr6.02, Bschr6.01 |
| 5 | Northeast Barents Sea (station 18 + 40) | Bschr10, Bschr3 |

#### Identification of a sex-differentiated region

A large ∼3 Mb (∼25–28 Mb) region on chromosome 5 was initially revealed by local PCA, defined by a two-cluster pattern. PCA of the SNPs within the region further confirmed that the two clusters corresponded to males and females separated along the PC1-axis and explained 34.1% of the variation (Figure 5). Calculations of *r*^2^ among all individuals revealed high levels of LD across the region (Figure S3), and the FST estimates between males and females revealed a significant genetic differentiation across the entire sex-differentiated region (Figure 5A and Figure S10). We further identified multiple genes annotated within this region with FST values between males and females ranging from 0.41 to 0.42, which included the *foxj3* gene and *phf20* like gene (which harbored SNPs that were tightly linked to sex in the polar cod in the present study) (Figure 5 and Table S7). The *phf20* gene was predicted as BOS_032872 by annotation and found to likely be PHD finger protein 20 (identities= 661/681(97%)) through Blast search. Inspections of the genotype composition of the region further uncovered similar patterns as using the sex-differentiating SNPs (see Material and Methods), with females generally being HOM REF and HOM ALT, and males were found to harbor more HET variants (Figure 5B).

**Figure 5:**
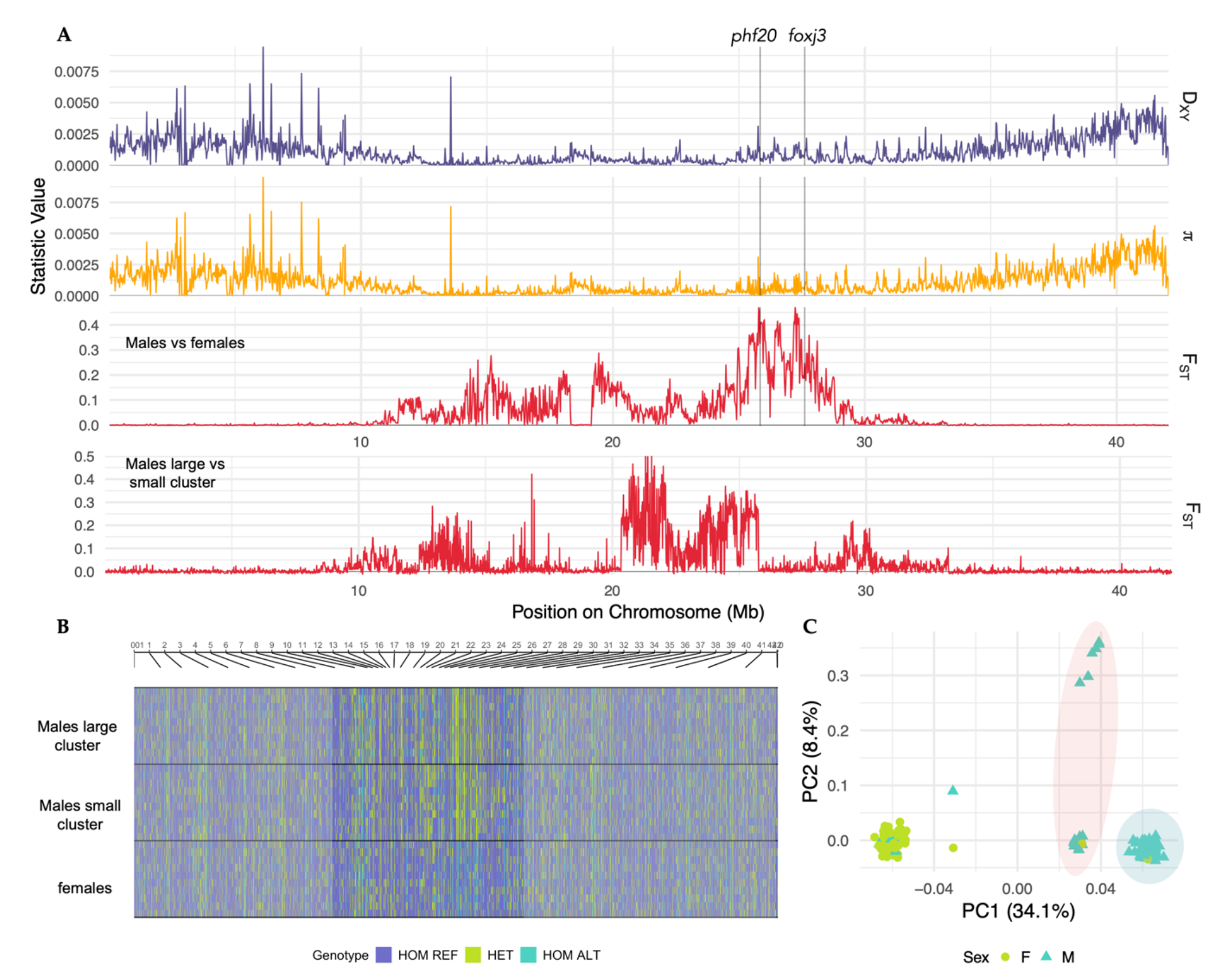
Large sexually differentiating region identified on chromosome 5 in polar cod. **A)** DXY, π, and FST estimates between males and females as well as between two identified male clusters along chromosome 5. A gene (*foxj3*) previously found to be involved in gonad development, and a gene found to harbor sex-linked SNPs (phf20) was identified within the region. **B)** SNP genotype composition of the sex region revealed males to be highly heterozygous across the region, compared to the female fish. **C)** PC1 plotted against PC2 of SNPs within the sex-determining region uncovered sub-structuring of male polar cod specimens, making up a large and a small cluster along PC1.

**Figure 6:**
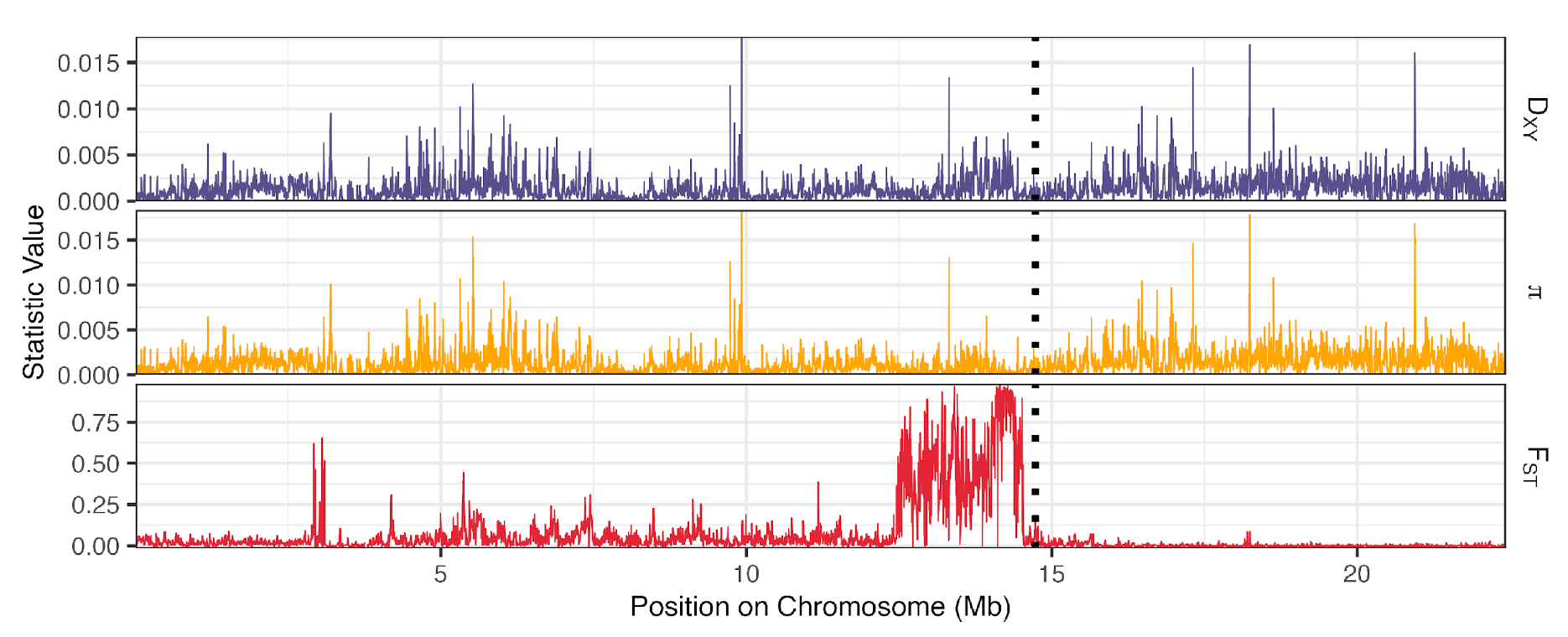
Genetic differentiation between homozygous haplotypes of inversion Bschr12.02 and genomic location of hemoglobin MN cluster. Estimates of DXY, average π and DXY between Hom common and Hom rare of inversion Bschr12.02. Chromosomal position of the MN cluster is marked with a stippled line.

Intriguingly, from the PCA plot (across the sex-linked region) we also detected a sub-structuring within males, where a smaller fraction of the males (n=20) separated into their own sub-cluster along the PC1-axis (irrespective of location; see Figure 5C and Figure S11). The FST estimates between these male-specific sub-clusters uncovered elevated estimates (ranging from ∼0.2-0.5) adjacent to and slightly overlapping (∼20- 27 Mb vs. ∼25–28 Mb) the main region differentiating between the sexes (Figure 5A and B). The biological importance of these regions was further confirmed by the selection analyses (Tajima’s D), uncovering that males, irrespective of cluster, experience balancing selection in the region (positive Tajima’s D in the region ∼19-30 Mb), whilst estimates for females seem to be more neutral (estimates close to 0 in the same region; see Figure S12).

## Discussion

By taking advantage of the newly generated polar cod chromosome-level genome assembly^44^ together with high coverage population sequencing data, we uncover a substantial number of chromosomal inversions as well as other genomic regions in high linkage disequilibrium (LD), which mainly constitute the five chromosomal fusions detected in polar cod^44^. The chromosomal inversions displayed high degree of LD (measured by *r*^2^) as well as genetic divergence (measured by FST) between the homokaryotypes. However, they varied notably in size, i.e., from smaller (∼3 Kb) to larger (∼15 Mb), where the largest was found to span almost half of the chromosome. Most of the identified inversions are seemingly *B. saida*-specific^44^, implying that they likely originated after the polar cod split with the ancestor of the gadids approximately 4-5 million years ago^13,49^. The observed differentiation within the inversions compared to surrounding chromosomal regions is concordant with what would be expected between homokaryotypes, due to low recombination in the heterokaryotypes^3,14,40,50^ . For some of the larger inversions (e.g., Bschr10, 13, 14, and 17) signatures of lower LD accompanied by lower FST estimates between the inversion homokaryotypes (occurring at the middle and/or towards the end of these inversions), indicate increased recombination, i.e., crossing-over events and gene flux^51–55^. Signs of increased recombination between the heterokaryotypes suggest that these inversions are relatively old^51,56,57^, supporting the hypothesis the inversion polymorphisms have been maintained within the species, and therefore are likely of ecological and evolutionary importance. Moreover, signals of selection were mainly detected in the breakpoint regions of the inversions, supporting the breakpoint mutation hypothesis^58^. This suggests that inversions originate and are selected due to adaptive mutations at breakpoint regions, where, for instance, adaptive variants are combined within or outside inversions depending on the orientation^58^.

The majority of the detected inversions display patterns of balanced polymorphism between the karyotypes (i.e., Bschr2, Bschr3, Bschr4.01, Bschr4.02, Bschr10, Bschr12.02, Bschr14 and Bschr16.01), which is suggested to be sustained by balancing selection^59^, hence also suggested by our selection analysis. Moreover, a few of the inversions show a skewed frequency pattern, with one of the homokaryotypes being more common and few heterokaryotypes being present, which indicates that these inversions could be under divergent selection^59^. However, there is also a possibility that the observed skewness is attributed to genetic drift. Additionally, some of the inversions seem to be partly linked, further supporting that these regions are of evolutionary importance, i.e., facilitating adaptation to specific environmental conditions and/or differences in life history traits. The phenotypic differences (reproduction and growth) of polar cod inhabiting Arctic vs. Atlantic fjords around Svalbard^31,60^, differences in phenotypic characteristics of polar cod in the Russian Arctic — such as size and color^61^, as well as the possible existence of local fjord populations^31–33^, point towards the presence of cryptic ecotypes with e.g., divergent migratory behavior and/or adaptation to different environmental conditions. Moreover, the presence of the complex inversion types — both those with a continuous divergence of one of the homokaryotypes (and the heterokaryotypes) as well as those with discrete clustering of three homokaryotypes and heterokaryotypes— suggest that these inversions are under divergent selection, with possibly high degree of gene flow between multiple ecotypes and/or sub-populations. However, these patterns could also be explained by secondary contact of historically isolated divergent sub-populations, as suggested by the three clades delineated by the mitochondrial phylogeny (see Figure S21 and previous studies^36^).

Detection of numerous chromosomal inversions as well as larger LD regions — including the chromosomal fusions and the sex differentiating region — indicate an evolutionary role of such genomic rearrangements. This hypothesis is further strengthened as the majority of these rearrangements were found polymorphic, and by such maintaining the genetic variation (of highly linked adaptive alleles) needed to rapidly respond to the extreme and variable environmental conditions that this cold-water fish species encounter. For instance, unlike the smaller sex-determining region identified in the closely related Atlantic cod^62,63^, the sex differential region in polar cod seems to be associated with one of the species-specific chromosomal fusion events detected (Hoff et al. submitted)^44^. This may be analogous to the chromosome fusions detected in two stickleback species generating a X1X2Y sex chromosome system^64,65^, suggested to be favorable by linking genes important for sexual dimorphism and mating behavior, and thus, promoting reproductive isolation^65,66^. Intriguingly, within the sex-differentiating region in polar cod, we identified several genes that harbored SNPs that clearly segregated between the sexes, including *foxj3* which have previously been found to display male specific expression patterns in spot-fin porcupinefish (*Diodon hystrix*)^67^. Furthermore, we also identified a narrower part of the sex-differentiating region which discriminates between two sub-groups of males. This male dichotomy could potentially be linked to ecotype differences due to mating behavior in polar cod (as found between stickleback species ^64–66^) and/or timing of reproduction.

For the functional aspect of the detected inversions, we highlight the two inversions that were found to be linked with the described hemoglobin gene clusters: the LA and MN clusters^48^. For Bschr15, the LA gene cluster was localized within the inversion, while the MN cluster was found localized just outside of Bschr12.02, in close proximity to the distal breakpoint. Hemoglobin is an essential oxygen-transporting protein for a variety of animals, including fish^68–70^. In codfishes, the copy number of hemoglobin genes could be linked to the environmental conditions that the species inhabit, i.e., with the highest numbers of hemoglobin genes in those species that are found in shallower waters, such as Atlantic cod^71^. Moreover, at the intraspecies level, the Atlantic cod is shown to harbor non-synonymous, tightly-linked SNP polymorphisms at the β1 locus^72,73^, thought to be associated with thermal adaptation^72,74^. Given that polar cod encounter a variety of environmental conditions, i.e., both occupying Atlantic and Arctic fjords around Svalbard and that they in early life stages can be typically found more associated with ice sheets than the older age classes^75–78^, it is likely that genetic polymorphisms with potential functional shifts in hemoglobin genes are favorable. For instance, having hemoglobin genes located within an inversion could allow different variants of the genes and/or regulatory elements to develop and be maintained among different subgroups of polar cod, in the presence of gene flow, hence enabling an extended repertoire of gene variants in the species.

When utilizing the inversions as genetic markers in genotype clustering analyses, a potential sub-structuring of the polar cod specimens within the Barents Sea and adjacent regions became apparent, which contrasts with the little or no divergence detected using the genome-wide SNP dataset (pruned for linkage and excluding the inversions). These findings are not surprising if we assume that the chromosomal inversions make up a genetic basis for ecotype divergence and/or local sub-population differences, as observed in Atlantic cod and other marine fish species^10,14,50,79^. Outlier loci or SNPs found to be associated with divergent selection between subgroups of a population can also reflect similar overall patterns as neutral markers^80^, or in some cases have the power to delineate a finer scale population sub-grouping than neutral markers^14,81–84^. However, it should be noted, that our sampling scheme is not designed to fully elaborate on cryptic ecotype or local sub-population differences, mainly due to the potential high degree of mixing of the different ecotypes and sub-populations at the feeding grounds, where most of the samples included in this study are collected from. We anticipate that a clearer structuring would become evident if we lowered the heterogeneity most likely present in our dataset.

To summarize, the detailed characterization of the genomic signatures of the keystone fish species in the Arctic — the polar cod — shed light on the potential importance of chromosomal rearrangements in local adaptation as well as sub- population structuring in this species, despite high degree of gene flow within the Barents Sea and adjacent regions. Maintaining such genomic structures and their variants is likely of high importance for enabling polar cod to tackle the forecasted reduction in ice sheet coverage and ocean warming.

## Materials and Methods

### Samples, DNA extraction, and genome sequencing

For the examination of population genomic differences in the Barents Sea, Svalbard, and adjacent areas, a total 290 polar cod individuals were selected for population-level whole genome sequencing (Tables S8-17). The samples were from different locations across the Barents Sea and adjacent regions, including the two hypothesized spawning grounds of the polar cod Northwest of the Barents Sea as well as Southeast, off the coast of Novaya Zemlya, Russia (Figure 1D and Table S8). Samples from Iceland, as well as a few samples from Tyrolerfjorden, Greenland, and Davis Strait, Canada were included. For the Barents Sea, we also included samples from different seasons and years to enable the detection of spatiotemporal and cohort genomic signatures (for more details see Supplementary Materials and Methods and Figure S13).

Genomic DNA was isolated from finclip for all samples except for the Northwest Barents Sea (gill rakes were used) and Hornsund samples (where spleen was used). The OMEGA Mag-Bind® Blood & Tissue DNA HDQ 96 Kit (Omega Bio- tek) kit was applied for isolation following the kit guidelines and performed on the KingFisher Flex 711 robot (Thermo Fisher Scientific). Sequencing libraries were prepared using KAPA Hyper kit (Roche) and genome sequencing was performed by the Norwegian Sequencing Centre (NSC; https://www.sequencing.uio.no), University of Oslo, Norway using the HiSeq4000 System (Illumina) in 2x150bp mode (150bp paired end) for more details see Supplementary Materials and Methods.

### Mapping reads, variant calling, and quality filtering

The sequencing data were mapped to the new chromosome-level polar cod reference genome^44^ and resulted in a median mean read depth of ∼18× across all individuals (Figure S14). Variant calling was performed with the Genome Analysis Toolkit (GATK)^85^ pipeline (for more information see Supplementary Material and Methods) and yielded a total of 25,633,802 raw SNPs, where mean individual SNP depth was calculated to range from 12× to 19× among individuals (Figure S15). Initial hard filtering of the SNP variant dataset was performed using the GATK “VariantFiltration” with filtering settings according to GATK recommendations: StrandOddsratio (SOR) > 3.0, FisherStrand (FS) > 60.0, MQ < 50.0, MappingQualityRankSumTest (MQRankSum) < -12.5, ReadPosRankSumTest (ReadPosRankSumneg) < -5.0.

Due to detection of an excess of heterozygous sites (with low quality score) – by manual inspection of the QualByDepth (QD) – we adjusted the QD filter setting to QD < 5.0 (see Supplementary Materials and Methods, Figure S16, S17 and S18). To ensure a panel of high-quality SNPs, we further filtered the dataset using VCFtools v0.1.16^86^ applying the following options and settings: MAF=0.01, max-missing=0.8, minQ=30, minDP=10, maxDP=50, min-meanDP=10, max-meanDP=50, max_allele=2, min_allele=2, mac=2. The quality filtering resulted in a complete set of ∼2.7 million high quality bi-allelic SNPs across the 18 chromosomes (Table S18).

### Delineating the population structure of polar cod in the northern Barents Sea

The presence of population genomic structure was investigated by principal component analysis (PCA) using PLINK v1.9^87^ and visualized using R v4.0.3^88^. An initial inspection of the quality-filtered SNP dataset, uncovered a distinct three-cluster pattern when plotting PC1 against PC2, explaining 7.57% and 6.51% of the variation, respectively, including all samples (n=290). All sample locations were represented across all three clusters (Figure 1C). The distinct three-cluster pattern is indicative of larger linked regions being present in the genome. For the other dimensions, i.e., PC2 vs. PC3 and PC3 vs. PC4, these signals (of linked regions) were not observed, and thus, all samples clustered together in one large group (Figure S2), indicative of no strong population structure in our dataset.

Moreover, based on the discoveries of multiple larger linked regions in the polar cod genome (see detailed description of the inversions and fusions below), we created a putatively neutral SNP dataset where highly linked regions of SNPs were removed. First, all regions identified to display high LD using BEDtools v2.30.0^89^ were excluded (Table S19). Second, the dataset was linkage pruned using BCFtools v1.12^90^ with the option “+prune” and the settings: -m 0.2 and -w 100kb, to exclude additional remaining linked SNPs. This resulted in a genome-wide putative neutral SNP set of a total of 564,970 SNPs distributed across the genome.

Subsequently, we re-ran the PCA analyses as well as ADMIXTURE v1.3.0^34^, using the above mentioned neutral SNP set. For the ADMIXTURE analyses, the number of divisions was run up to K = 10, with cross-validation (CV) estimates ranging from 0.30545 for K=1 to 0.39216 for K=10 (Figure S19). Based on the CV estimates, which were rather low, we here present the most informative ancestry fractions for ADMIXTURE using K=3. The ADMIXTURE ratios were visualized using R v4.0.3^88^. Mitochondrial phylogeny and demographic analyses were also conducted (for more information see Supplementary Material and Methods, Figure S20 and S21).

### Detection of chromosomal inversions and fusions

#### Local PCA with lostruct

To explore the genetic variation across the genomes of our 290 polar cod free of assumptions of any population structure or clusters, we ran the R package lostruct^91^ in windows of 100 SNPs along the chromosomes. Lostruct calculates PCA in non-overlapping windows along the chromosomes and visualizes the variation among the individuals on a local level by multidimensional scaling. The method allows for a nuanced view of the variation among individuals locally along the chromosomes. All regions displaying high variation among individuals were extracted for manual inspection based on location on the chromosome.

#### Linkage Disequilibrium along chromosomes

To estimate linkage between SNPs along the chromosomes of polar cod, PLINK v1.9^87^ was used to remove sites with more than 0.01% missing data and SNPs were randomly thinned down to 10% of the original count. After that, PLINK v1.9^87^ was used to calculate *r*^2^ via the option --r2 across all 18 chromosomes, within the 290 individuals. Visualization of *r*^2^ was done in R v4.3.2^88^ as a heatmap using the R package Scattermore^92^.

#### Identification of chromosomal inversions

The SNPs located in the regions displaying signals of high MDS variation among individuals accompanied with a three-cluster pattern on the initial lostruct analysis were extracted for each of the putative inversions, separately. For these putative inversions we performed the following steps to confirm if the variation observed followed the expected signals of a chromosomal inversion: 1) A PCA was performed on SNPs within the localized region reported, to investigate discreteness of clustering between individuals. 2) Heterozygosity was calculated across the extracted region and PC1 vs PC2 plots were colored to display individual levels of heterozygosity for the given region to investigate if the middle cluster displayed a higher level of heterozygosity, as would be expected from an inversion. For chromosomal inversion showing discrete clustering, clusters were determined both by using a k-means based approach using the find.clusters() part of the “Adegenet” package^93^, as well as a visual approach where heterozygosity was taken into account. The individuals in each cluster were assigned to Hom rare: the homozygous cluster including fewest individuals, Het: the heterozygous middle cluster, and Hom common: the homozygous cluster containing the majority of individuals. For chromosomal inversions displaying divergent homozygous types, and non-discrete clustering between genotypes, find.clusters() was used to determine clusters. The inversions identified were categorized as either i) classical displaying three discrete clusters, or ii) complex displaying additional sub-structuring and/or continuous distribution of samples between clusters. 3) *r*^2^ was calculated across all 18 chromosomes, additionally for the discrete inversions, *r*^2^ was calculated across the chromosome among all the individuals, and only among individuals harboring the Hom common variant, the expectation being that linkage will be much higher when comparing all individuals, than among individuals that are homozygous for the inversion, across the region of which the inversion is located^1^.

Moreover, we characterized the inversion homokaryotypes differentiation for all the classical inversions as well as for a selection of the divergent inversions, by estimating FST, DXY and π using Pixy v1.2.6^94^. For the classical inversion, this was conducted between individuals that were defined in the different homozygous clusters, i.e., the Hom common vs. Hom rare, while for the divergent inversions between individuals defined in the hom1, hom2 and hom3 clusters. For the divergent inversions, estimates were calculated for Bschr6.02, Bschr7 and Bschr16.02 of which we had sufficient numbers of specimens of each homozygous genotype cluster. Neighbor-joining (NJ) trees based on polymorphism were constructed for inversions displaying three genotypes as well as one complex inversion (Bschr16.02), to further validate the observed clustering of individuals using the R package “Phangorn”^95^. In this analysis, alleles of heterozygous sites were randomly sampled to create one pseudo-sequence per individual before inferring the NJ tree.

For all identified inversions that displayed a three-genotype pattern, frequencies of genotypes were visualized as pie charts with R using ggplot2^96^ and grouped across sampling locations to investigate distribution pattern.

Lastly, for a selection of the inversions we investigated the SNP genotype calling (classified as HOM ALT, HOM REF or HET in accordance with the reference genome) along the chromosomes by visualizing the genotype distribution using the R package “GenotypePlot”^97^.

#### Correlation between inversions

For the inversions displaying three genotypes, we tested for association between paired inversions by conducting an inversion correlation analysis (using Pearson’s product moment correlation with the cor.test() function in R).

#### Selection analyses

Signals of selection across the entire genome of the polar cod were conducted by computing Tajima’s D in windows of 25 Kb using VCFtools v0.1.16^86^.Tajima’s D was calculated per location, as well as per sampling year and season for the Northwest Barents Sea samples.

Further, a phased dataset was generated to enable additional analyses of iHS and XP-EHH. For this, the VCFs were phased using Shapeit4 v4.1^98^ chromosome by chromosome, with settings --window 0.5 -T 4. Estimates of iHS and XP-EHH were obtained by using the R package “rehh”^99^. IHS was calculated across each sample location individually, by first using the function data2haplohh with option polarize_vcf=FALSE, then scanned using the function scan_hh, and lastly iHS was estimated using the function ihh2ihs. XP-EHH was calculated pairwise for pairs of locations between Barents Sea Aug 2018, Billefjorden, and Russian samples, in total 30 comparisons across each chromosome. Calculations were done by first reading data using data2haplohh() with option polarize_vcf=FALSE, then using pop_scan() for each of two sample locations, and lastly pairwise XP-EHH calculation was performed using ies2xpehh(). All visualization was done in R using ggplot2.

#### Characterization of a sex-differentiating region

Upon running lostruct, we discovered a large region on chromosome 5 clustering our samples in two groups. We extracted variants across this region, ran and plotted PCA against metadata information on sex determined by morphology. Additionally, to validate the sex-linked region and its boundaries, we ran FST between the males and females determined from morphology using Pixy v1.2.6^94^ in 25 Kb windows, whilst between male sub-clusters 10 Kb window was used. To identify potential genes that might be involved directly in sex determination and/or gonad development within this region, we extracted genes from the annotation file overlapping with FST windows with values > 0.4 between the sexes. We identified linked SNPs within multiple genes extracted manually, where the SNP variant status corresponded to heterozygous and homozygous for male and female respectively when compared to the individuals that were morphologically sex determined during sampling. Genes in which the SNP resided were determined by the annotation, or by using Blastn online Suite^100^ to search for gene matches where the annotation was not complete (i.e., hypothetical protein identified but gene not determined). Search was done using the options “standard databases” and “highly similar sequences”. Subsequently, we used these sex-differentiating SNPs to genotype the sex of the remaining polar cod individuals that were not sexed in our dataset, for instance the younger juveniles that had underdeveloped gonads that could not be sexed visually. We identified that our dataset is made up of a total 150 males and 140 females (Table S7), for which we re-ran the pairwise FST using Pixy v1.2.6^94^ in 25 Kb windows.

#### Inspection of the hemoglobin gene clusters

The two hemoglobin gene clusters, LA and MN, were selected for further inspection as these genes are well characterized within the Gadiformes^71^ lineage. The hemoglobin genes and flanking genes for Atlantic cod were collected from Baalsrud et al.^71^ and were queried against the reference genome of polar cod using BLAST 2.13.0+ ^101^ blastn tool. Hits were double checked against the annotation GFF3 file of the reference genome in IGV to confirm a match.

#### Population genetic divergence and diversity

Population genetic diversity was estimated by calculating DXY, average π as well as FST (Weir and Cockerham 1984)^35^ along the genome in widows of 25 Kb within and between sample locations using Pixy v1.2.6^94^. Estimates were performed for our main locations: Northwest Barents Sea (summer 2018), Billefjorden, the 0-group collected in summer 2018, and Northeast Barents Sea (combining all five stations along the coast of Novaya Zemlya) against all other locations. In addition, estimations were done between specimens collected from the Kara Sea (station 9) as well as northern Novaya Zemlya (station 18) against the other Russian stations (as well as all other locations) to evaluate levels of differentiation within the northeast Barents Sea (not shown). Furthermore, we calculated a proxy for a global pairwise FST between sample localities by averaging the estimated values obtained by FST across all chromosomes for above mentioned comparisons.

#### Using chromosomal inversions as genetic markers in exploration of sub-clusters

Using the k-means method with function kmeans() in R, we conducted a simple clustering analysis on the genotype frequencies of 19 inversions to group the observations by genotype. First, we analyzed the 5 East Barents Sea sampling locations independently, grouping them in k=1 to 4 clusters (1 meaning no difference), and found that they could be grouped in three different clusters. In a second step, we conducted a new clustering analysis with the remaining sample locations and the three East Barents Sea clusters, sorting the sample in k=1 to 6 clusters, and found that they could be grouped in 5 different clusters.

#### Chromosomal inversions and Hardy Weinberg Equilibrium

Tests for Hardy Weinberg Equilibrium (HWE) of genotype frequencies among the 290 polar cod collectively, as well as within the five subclusters separately, were performed on biallelic genotyped inversions using the R package “HardyWeinberg”^102^. Moreover, FIS was estimated by using the function summary () from the R package “Adegenet”^93^ and calculated by (genind_summary$Hexp- genind_summary$Hobs)/genind_summary$Hexp, to determine if deviations from HWE were observed, could be attributed to heterozygote excess or deficiency.

## Supporting information

Supp_population_genomics_boreogadus_saida

## Acknowledgments

Library preparations and sequencing were performed by the Norwegian Sequencing Centre, University of Oslo. The computations were performed on resources provided by Sigma2 - the National Infrastructure for High Performance Computing and Data Storage in Norway. We thank Anna H. Ólafsdóttir and the scientific crew of the Árni Friðriksson (MFRI) for assisting on the sampling of polar cod specimens during the IESSNS surveys. We thank Leif Christian Stige (CEES, University of Oslo) for aiding in the sampling of polar cod in the Northwest Barents Sea. We thank Alexei Bambulyak, Vladimir Savinov (Akvaplan-niva) and Evgeniia Raskhozheva (Murmansk Marine Biological Institute, Murmansk, Russia) for assisting on the sampling of polar cod specimens from the Northeast Barents Sea. We also thank Julie Bitz-Thorsen for aiding in sampling in the Northeast Barents Sea. Lastly, we thank Alexandra Viertler for the polar cod illustration. We thank the crew of RV Kronprins Haakon for facilitating the trawling and sample acquisition in the rough Barents Sea.

## Funding

This work was funded by the Research Council of Norway through the following projects: **‘Nansen Legacy’** (RCN no. 276730) and the **ComparaCod** project (RCN no. 222378). Parts of the project were also funded by **MEMO-PRO** project (ID: RFMEFI61616X0073), the Murmansk Marine Biological Institute.

## Author contributions

SJ and SNKH conceptualized the study. SNKH and MFM isolated DNA for sequencing, handled and processed the data, as well as performed the following analyses: MFM mapped sequence data and performed the variant discovery pipeline, contributed to building scripts for analyses and visualization, and conducted the female estimation of Ne and mitochondrial phylogeny. SNKH performed variant quality filtering and performed downstream analyses including assessment of population genetic structures, selection analyses, estimating population genetic diversity, chromosomal inversion discovery and characterization, identification of sex region, estimating LD, characterization of sex associated region, and identification of hemoglobin regions. JMD performed sub-cluster analyses using inversions, tested for association between inversions, as well as estimated age for specimens where not recorded. MR created and provided scripts, aided in variant discovery pipeline as well as downstream analyses. ALM created and provided scripts as well as aided in analyses of chromosomal inversions. JAG coordinated the otolith age reading. Following co-authors sampled and provided specimens for sequencing from Northeast Barents Sea: PER, DM and ER; Northwest Barents Sea: SJ and SNKH; Iceland: CP; Greenland: KP; Kongsfjorden: HH and FC; Billefjorden and Hornsund: JN and IV; and Davis Strait, Canada: IRB. OKT aided in data management. Funding acquisition by SJ, KP, PER and KSJ. Visualization and design of figures done by SNKH, MFM and SJ. SNKH and SJ wrote the original manuscript, with relevant sections contributed by MFM and JMD. All co-authors read, provided feedback and improved the manuscript.

## Notes

### Competing Interest Statement

The authors have declared no competing interest.

