## Supplementary material for "Population divergence manifested by genomic rearrangements in a keystone Arctic species with high gene flow": Supp_population_genomics_boreogadus_saida

#### Supplementary html files

Supplementary html: classical inversions
https://www.mn.uio.no/ibv/personer/vit/snhoff/html_classical_inversions2.html

Supplementary html: FST
https://www.mn.uio.no/ibv/personer/vit/snhoff/html_fst_dxy_pixy_pixy.html

Supplementary html: Tajima's D
https://www.mn.uio.no/ibv/personer/vit/snhoff/html_tajimasd_final.html

Supplementary html: iHS
https://www.mn.uio.no/ibv/personer/vit/snhoff/html_ihs.html

Supplementary html: XP-EHH
https://www.mn.uio.no/ibv/personer/vit/snhoff/html_xpehh.html

Overview page for online Supplementary Materials:
https://www.mn.uio.no/ibv/personer/vit/snhoff/phd-files.html

### Supplementary Notes and Figures

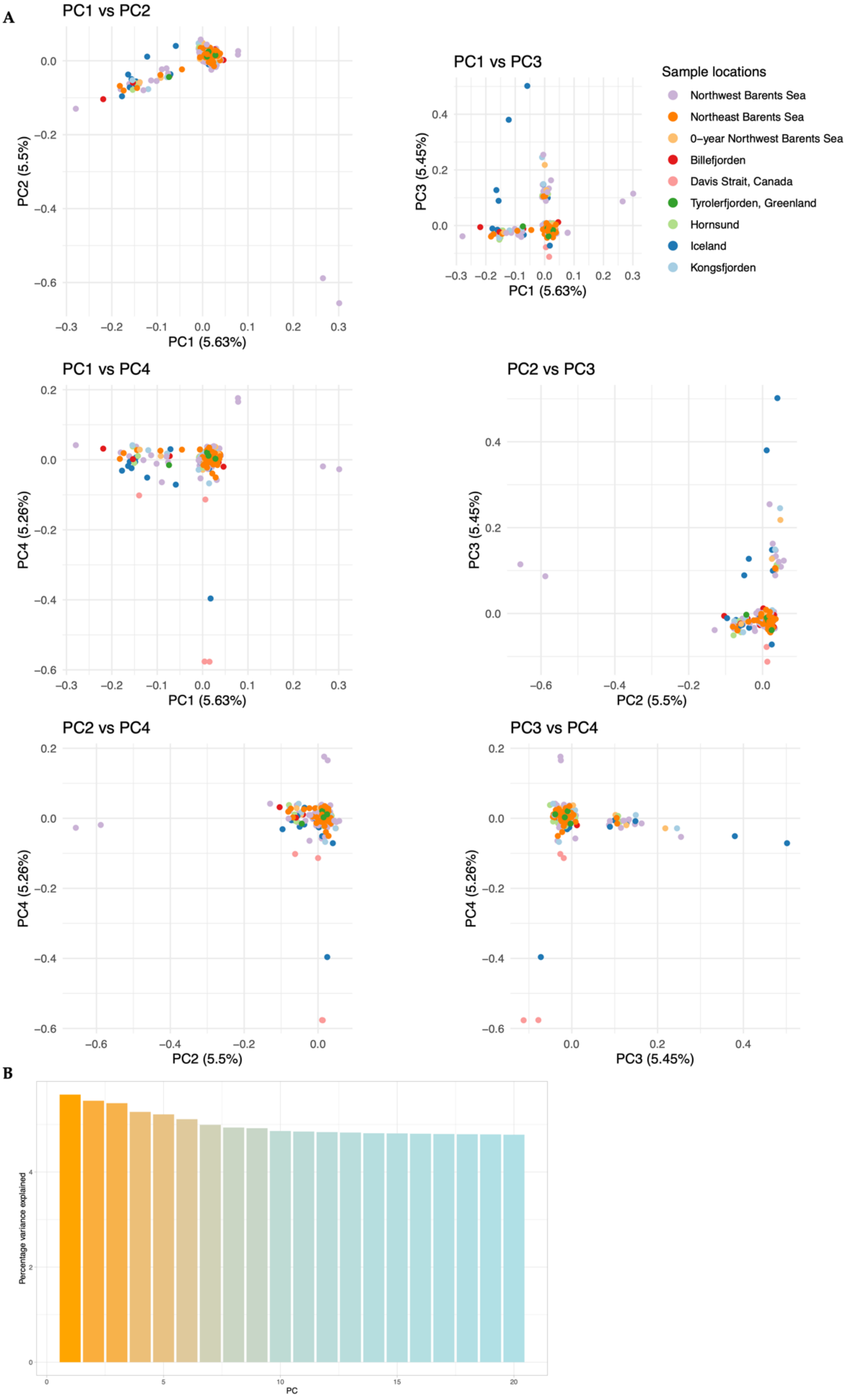

**Figure S1. A)** Principal component analyses across genome-wide high-quality SNPs posterior removal of linked regions and pruning (n=564 970). One tight cluster including samples from all geographic locations is observed while inspecting PC1 vs PC2. A tailing of specimens can also be seen along PC1, likely because of some.
**B)** Percentage of PC variation explained PC1 through PC20.

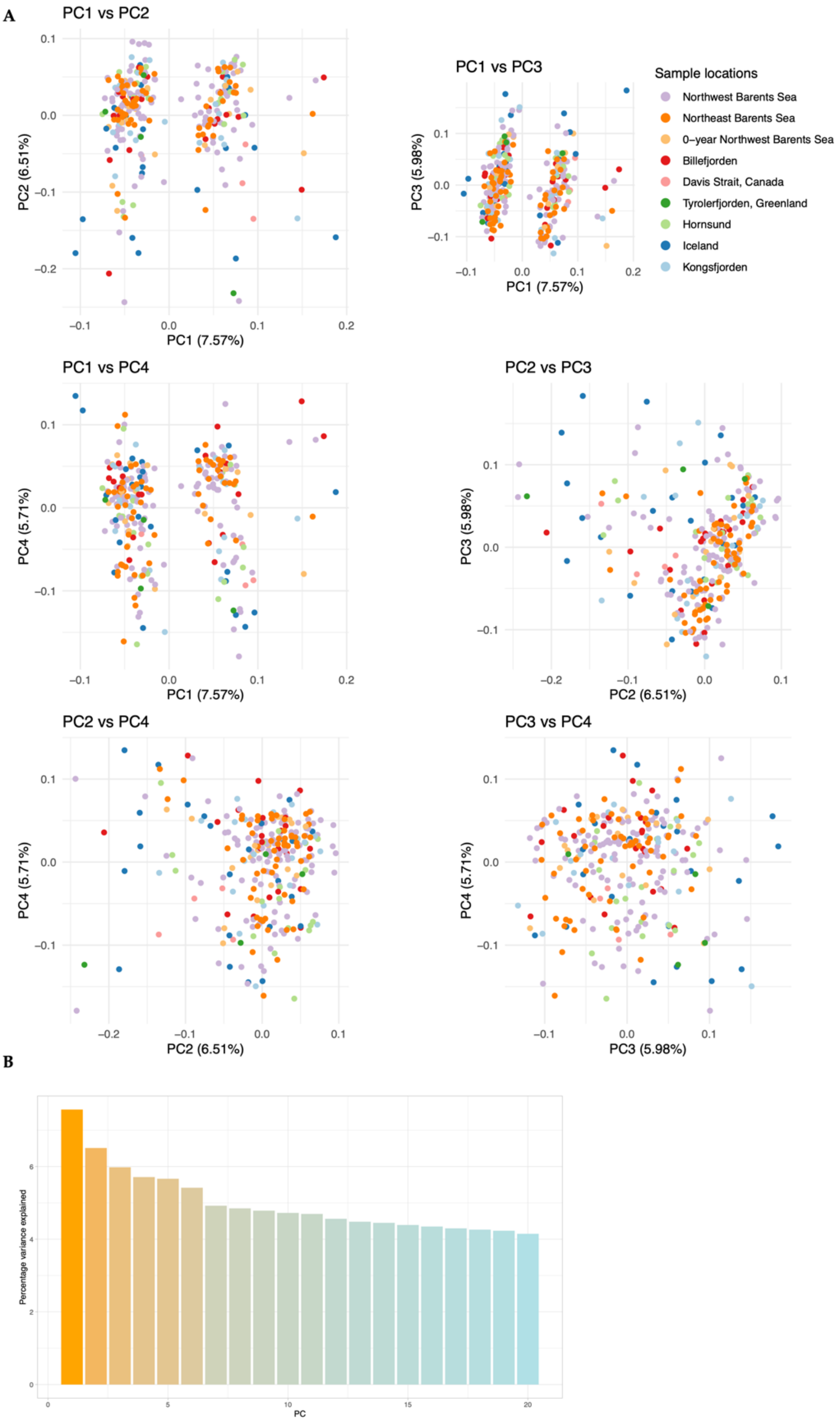

**Figure S2. A)** Principal component analyses across genome-wide high-quality SNPs (n=2 705 025) prior to removal of linked regions and pruning. A three-cluster patterning among samples can be observed along PC1, resulting from polymorphic structural variants across the genome of polar cod. **B)** Percentage of PC variation explained PC1 through PC20.

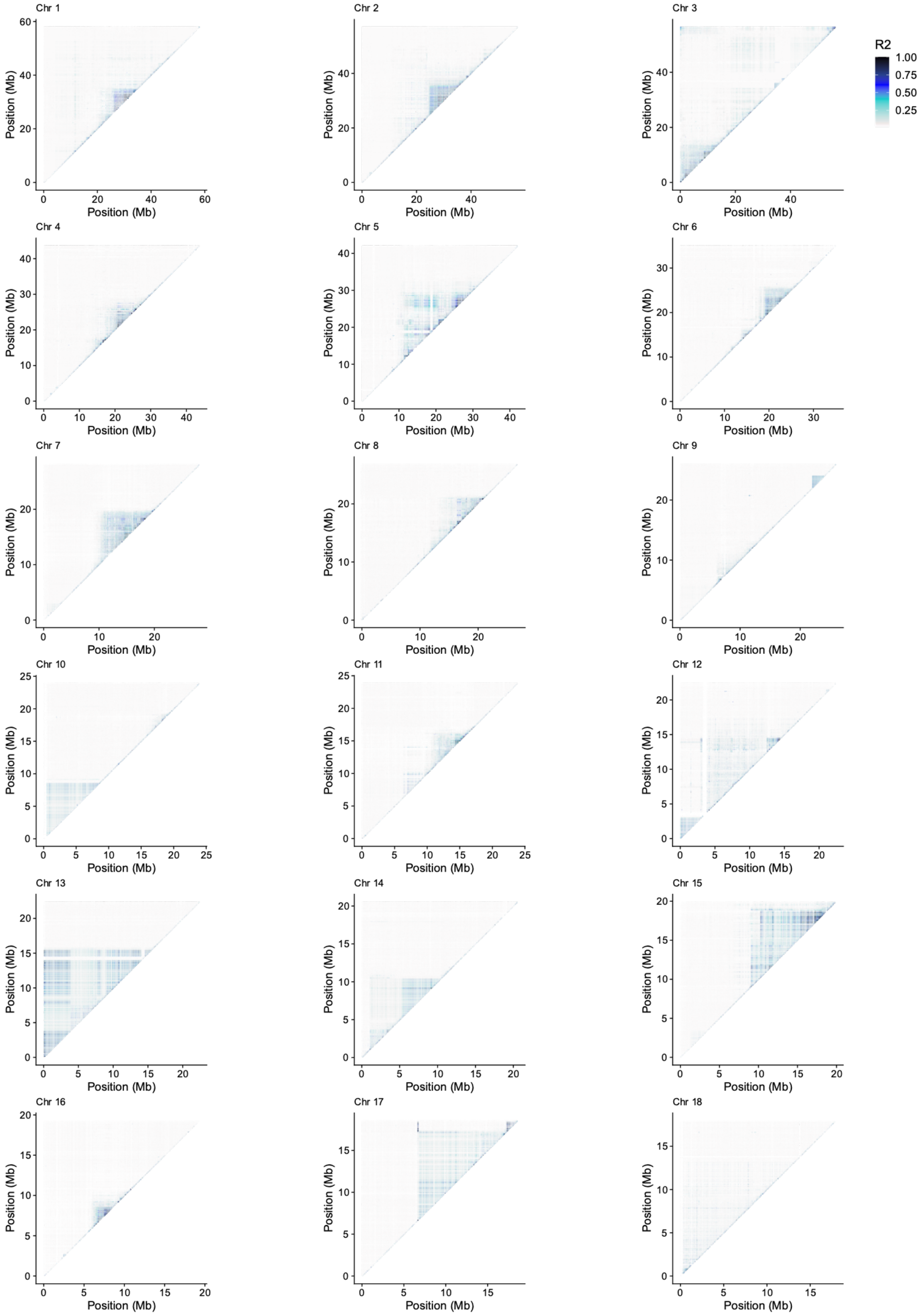

**Figure S3.** Estimates of linkage disequilibrium (measured as $r^2$) are shown as a heatmap among all 290 specimens along all chromosomes. Elevated estimates of $r^2$ among the samples were observed across many regions of the genome.

#### Supplementary Note 1

For the 11 classical inversions (Table S2) the PCAs of SNPs within these inversions displayed discrete three-cluster patterns with PC1 explaining from 27.6% and up to 71.9% of the variation observed (Figure 3A-G and Supplementary html: classical inversions). Evaluation of heterozygosity levels of the three discrete clusters revealed substantially higher heterozygosity of the middle cluster, compared to lower heterozygosity of the two flanking clusters (Figure 3E and Supplementary html: classical inversions). The difference in heterozygosity of the clusters is in line with what would be expected from chromosomal inversions where the two lower heterozygosity clusters each represent individuals carrying the homozygous haplotypes, and the middle cluster represents the individuals that are heterozygous for the inversion. We proceed to genotype the clusters accordingly (Figure 3E and Supplementary html: classical inversions). Neighbor-joining (NJ) clustering analyses of SNPs within the inversion region further validated that the samples clustered according to the genotypes assigned to them (see Figure 3D, not shown for the rest of the classical inversions).

Of the total 20 inversions identified, nine inversions were classified as «complex» (Table S2). These inversions were found to display different levels of divergence for one of the two homokaryotypes (resulting in three homokaryotype clusters). Bschr6.02, Bschr7, and Bschr16.02 are the most extreme cases, found to exhibit six more or less discrete subclusters on PCA of SNPs within the inversion region (Figure 4C, S4B, and S5B). By visualizing heterozygous sites (%) overlayed on PCA we found that the individuals making up the three corner clusters (PC1 vs PC2) were highly homozygous in comparison to the three centrally located, which displayed significantly higher levels of heterozygosity (Figure 4E, S4B and S5B). NJ clustering of SNPs within the region for Bschr16.02 resulted in subgroups corresponding to the subgroups delineated by PCA (Figures 4C and D). With Hom1 and Hom2 grouping together, and Het2 located between them. Furthermore, Hom3 was found located at the other distal end, with Het1 and Het3 in-between (Figure 4C and D).

Four of the larger inversions identified Bschr10, Bschr13, Bschr14, Bschr15, and Bschr17 were found to make up close to one half of a chromosome, with one of the boundaries located in the distal end of the chromosome and the second close to the middle of the chromosome. Many of the smaller inversions were found to be located close to the ends, or centrally on the chromosomes (Figure 2A). Intriguingly, one of the complex inversions identified, Bschr2, was found located centrally on chromosome 2, overlapping with a region that makes up a fusion region between two ancestral chromosomes[1]. Collectively, the chromosomal inversions identified were found to contain from 29 genes in Bschr04.01 and up to 974 annotated genes in the largest inversion, Bschr13 (Table S2).

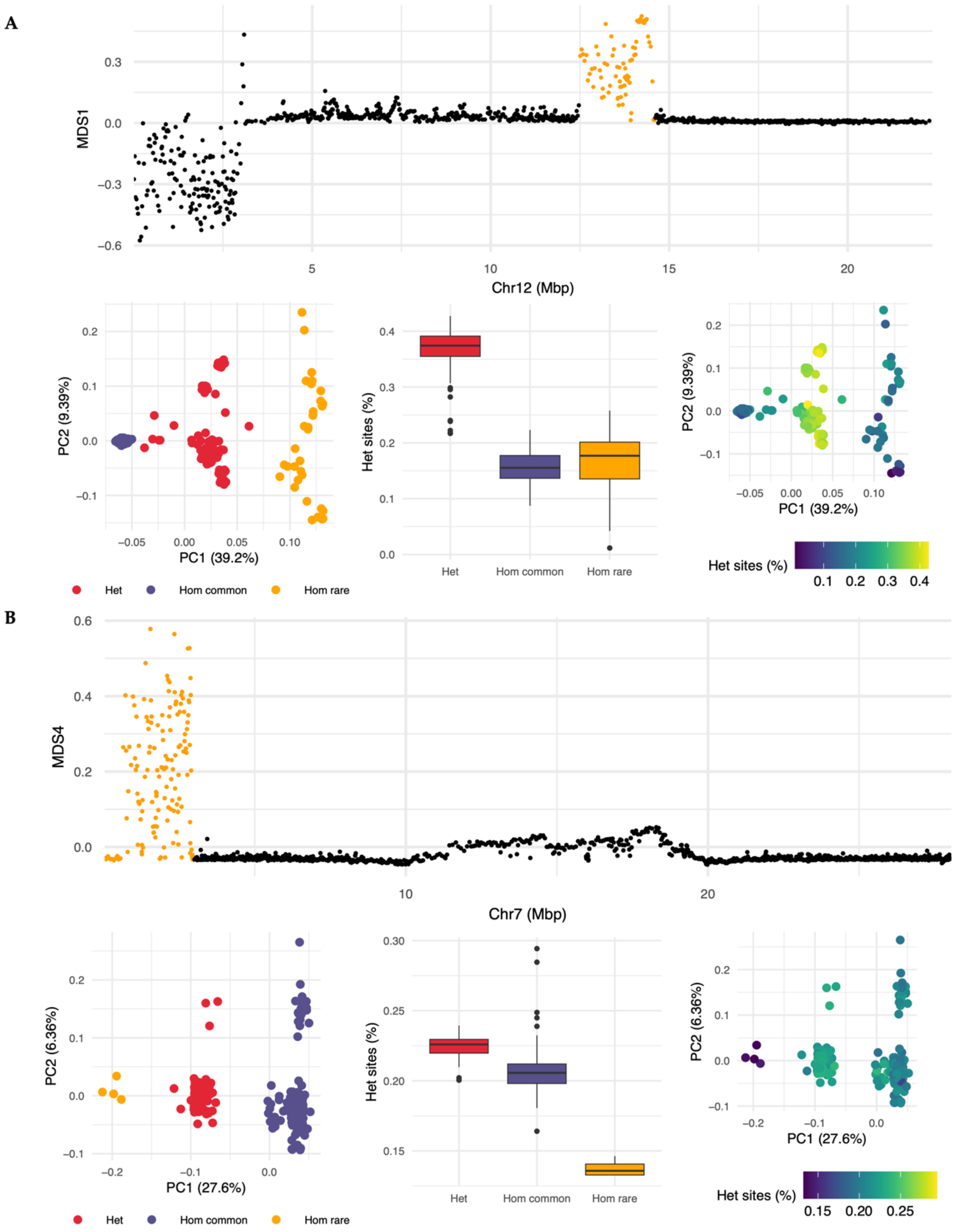

**Figure S4.** Identification and characteristics of two complex inversions where homokaryotypes (of one arrangement) as well as heterokaryotypes displayed a **A)** continuous (Bschr12.02) and **B)** dichotomous (Bschr7.01) pattern of divergence along PC2. Upper panel: inversion region shown in yellow. Bottom left: Principal component analysis across SNPs in the identified region. Bottom center and right: heterozygosity visualized across the three subgroups.

**Inversion Bschr6.02**

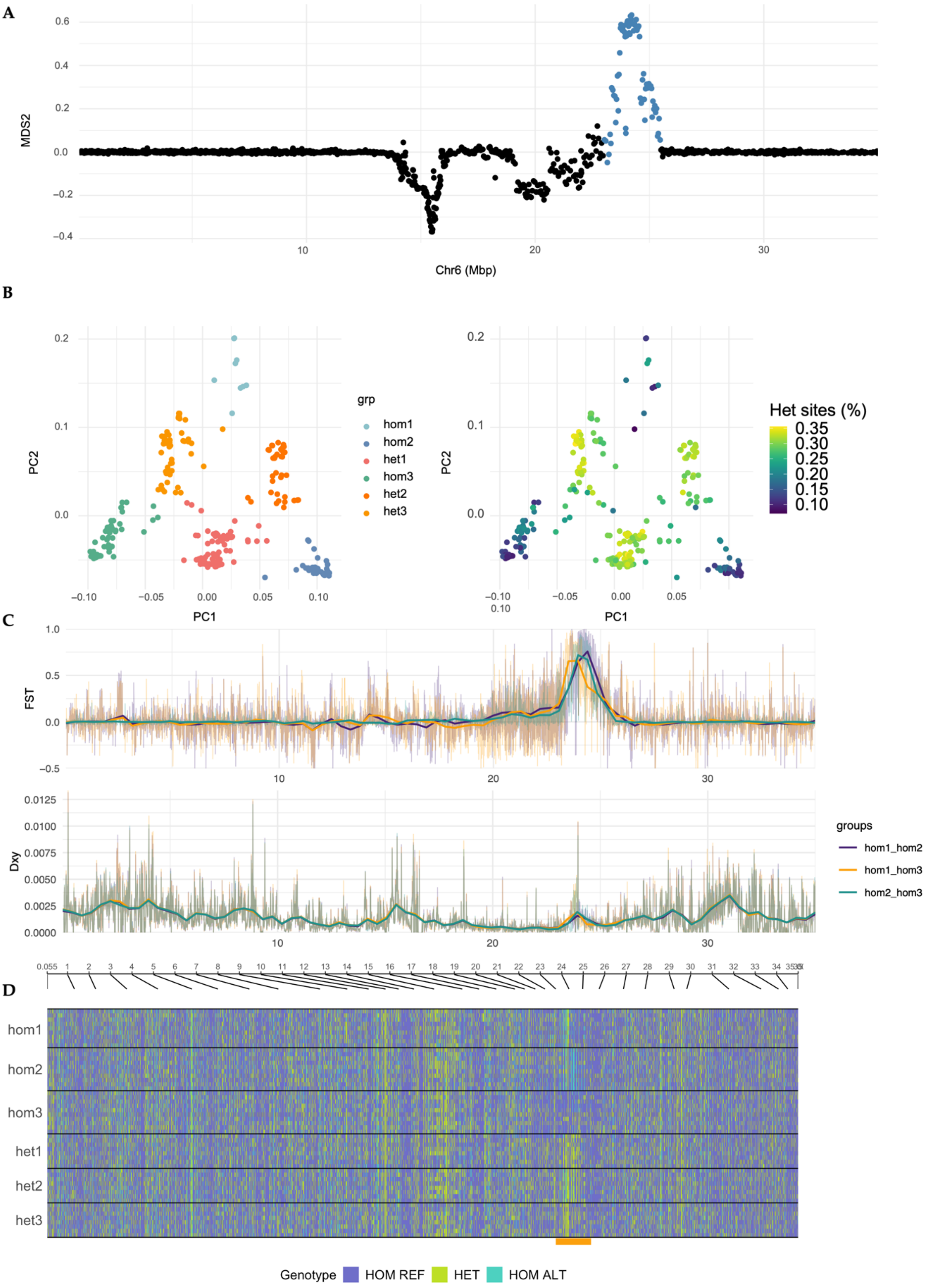

**Figure S5.** Identification and characteristics of complex inversion Bschr6.02. **A)** MDS1 variation along chromosome 6 generated by lostruct[2]. **B)** Left: PCA of SNPs residing within the inversion region. Right: PCA of SNPs residing within the inversion region with statistics for Het sites (%) visualized as a color scale. **C)**

Pairwise $F_{ST}$ and $D_{XY}$ (in 25Kb windows along chromosome 6) comparisons between individuals representing each homozygous cluster. **D)** SNP genotype composition along the chromosome (plotted using genotype_plot[3]), of a selection of samples of each genotype determined (i.e., being classified as either HOM ALT, HOM REF, or heterozygous (HET) in accordance with the reference genome), with the inversion location marked in yellow.

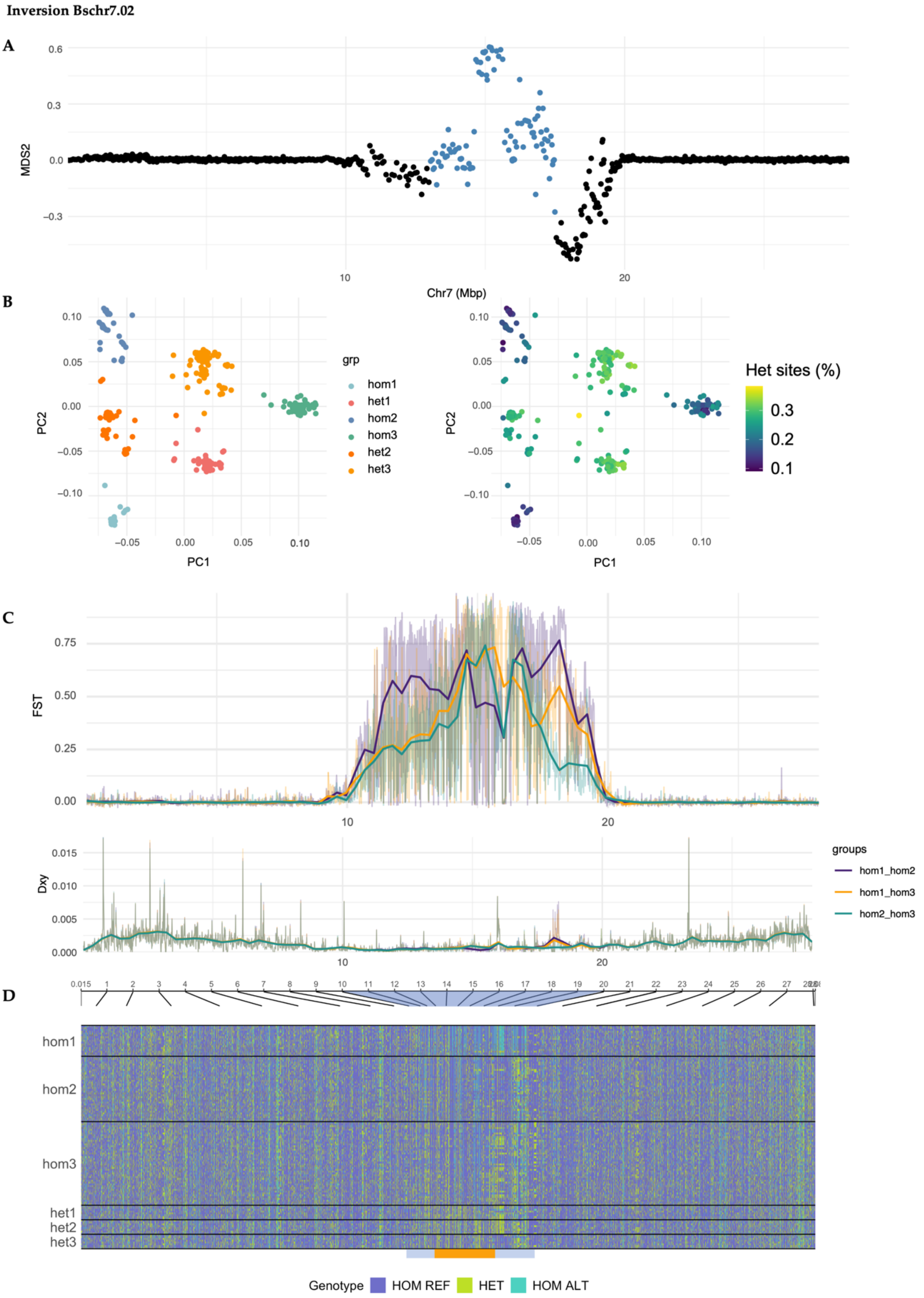

**Figure S6.** Identification and characteristics of complex inversion Bschr7.02. **A)** MDS1 variation along chromosome 7 generated by lostruct[2]. **B)** Left: PCA of SNPs residing within the inversion region. Right: PCA of SNPs residing within the inversion region with statistics for Het sites (%) visualized as a color scale. **C)**

Pairwise $F_{ST}$ and $D_{XY}$ (in 25Kb windows along chromosome 7) comparisons between individuals representing each homozygous cluster. **D)** SNP genotype composition along the chromosome (plotted using genotype_plot[3]), of a selection of samples of each genotype determined (i.e., being classified as either HOM ALT, HOM REF, or heterozygous (HET) in accordance with the reference genome), with the inversion location marked in yellow.

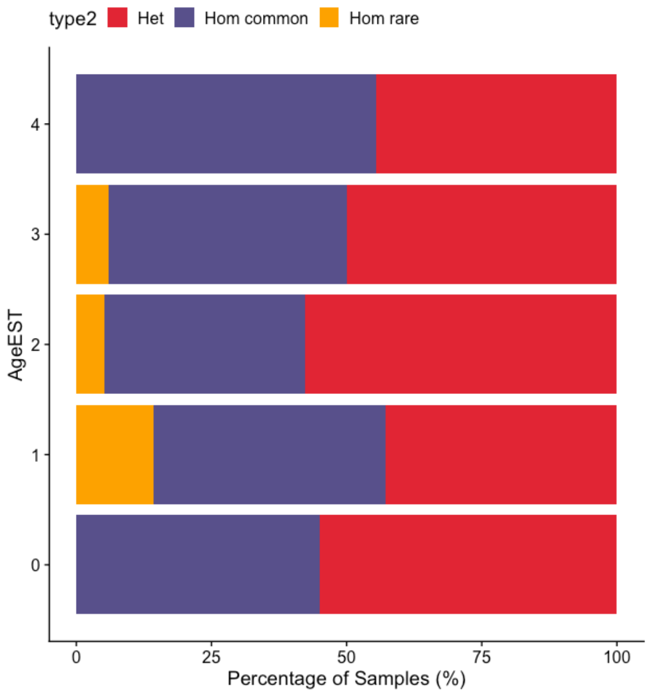

**S7.** Frequency of inversion Bschr14 genotypes in age classes. Y-axis show age of specimens in years. Colors denote inversion genotype.

#### Supplementary Note 2

Estimates of selection by the integrated haplotype score[4] (iHS) revealed smaller and larger stretches of positive or negative estimates within the chromosomal regions of the inversions, which were in line with some of the signals of selection detected by the Tajima's D analyses. For instance, for all locations of the inversion identified on chromosome 2 (~20 – 40 Mb) which overlaps with the chromosomal fusion, we detected an overall trend of elevated iHS estimates compared to surrounding regions (see Supplementary html: iHS). Similar patterns of elevated iHS estimates were detected for Bschr04.01, Bschr07.02. Moreover, positive estimates of iHS were found in parts of the inversions Bschr17 (central parts of inversion), Bschr16.02 (central parts of inversion), Bschr13 (latter part of inversion), and Bschr18 (first part of inversion) (Supplementary html: iHS). Inversion Bschr7.01 and Bschr04.02 are examples of inversion showing negative estimates of iHS within the inversion regions (Supplementary html: iHS).

When calculating the average nucleotide diversity ($\pi$) for the homokaryotypes for all of the classical inversions separately, we uncovered that for Bschr3 and Bschr4.02 the estimates were highly similar between the Hom common and Hom rare, whereas for Bschr4.01, Bschr5, Bschr12.01, Bschr13, Bschr14, Bschr16 and Bschr17 the estimates were highest for Hom common (Figure S9). The estimates for Bschr9 were found to be highest for Hom rare (Figure S9). For Bschr10 the estimates were comparable between the homokaryotypes for the first part of the inversion, while higher estimates for Hom common at the distal part of the inversion (Figure S9).
Estimates of cross-population extended haplotype homozygosity[5] (XP-EHH) between a selection of sampling sites (Northwest Barents Sea summer 2018, Billefjorden, and Northeast Barents Sea autumn 2019 vs. all other sites) revealed – in concordance with previous selection analyses – stretches or highly positive and/or negative values within inversion regions (Supplementary html: XP-EHH). For Bschr10 comparisons between Northeast Barents Sea autumn 2019 and Billefjorden, as well as Hornsund resulted in multiple dips of XP-EHH estimates (< -4) within the region of the inversions (see Supplementary html: XP-EHH). For Bschr9, we found that comparisons between Billefjorden and other locations revealed elevated estimates within the inversion, most likely attributed to inversion allele skewness (Supplementary html: XP-EHH). For Bschr4.02 comparisons between Northwest Barents Sea summer 2018 and most other sample sites revealed highly positive estimates of XP-EHH (Supplementary html: XP-EHH). For one of the larger inversions, Bschr13 local stretches (especially ~10-14 Mb) of positive values (>2.5) were detected within the inversion in comparisons between Northwest Barents Sea summer 2018 vs Barents Sea winter 2019, Hornsund, Kongsfjorden, Northeast Barents Sea autumn 2019 and the 0-group (Supplementary html: XP-EHH).

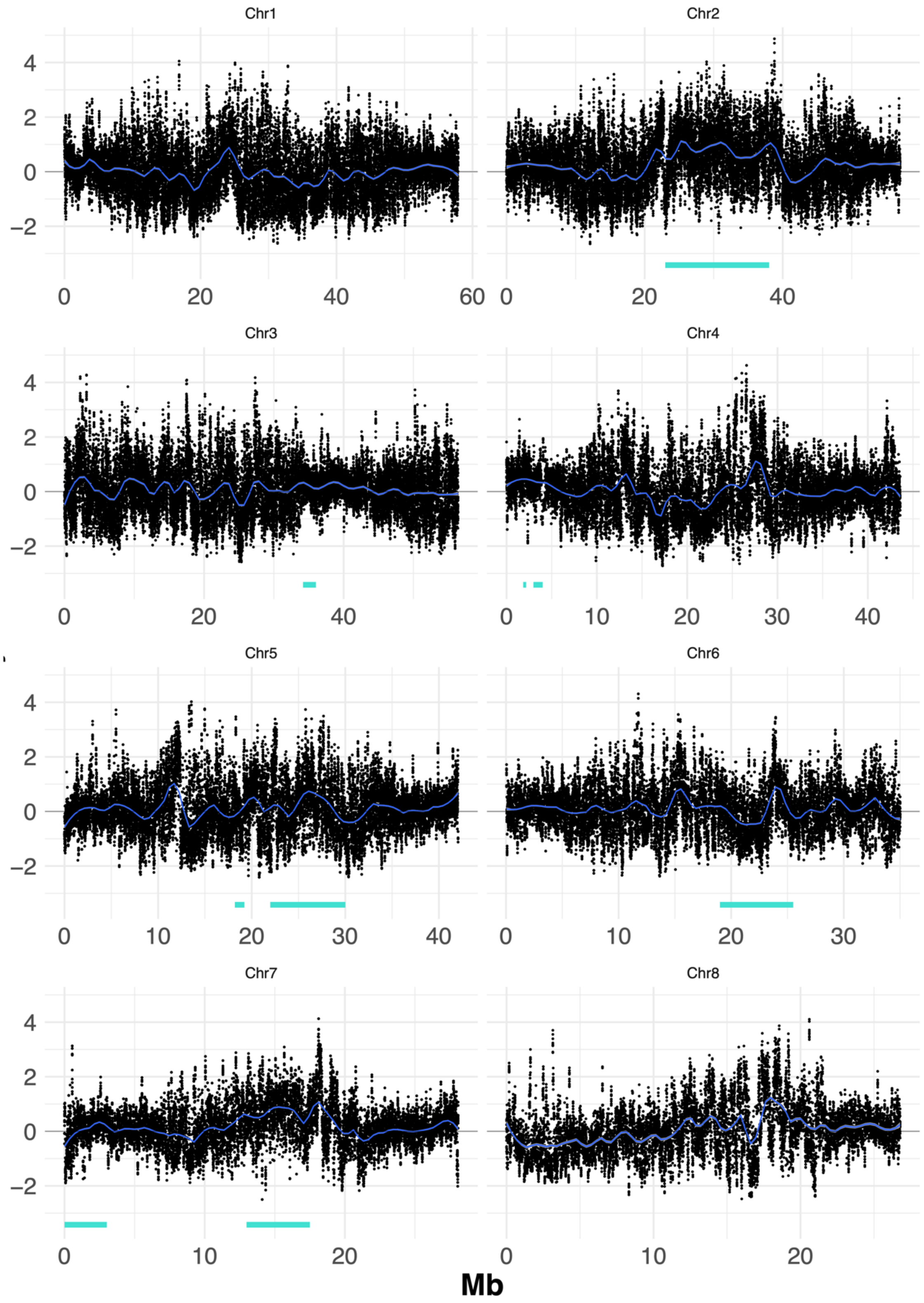
Chr1
Chr2
Chr3
Chr4
Chr5
Chr6
Chr7
Chr8
Mb

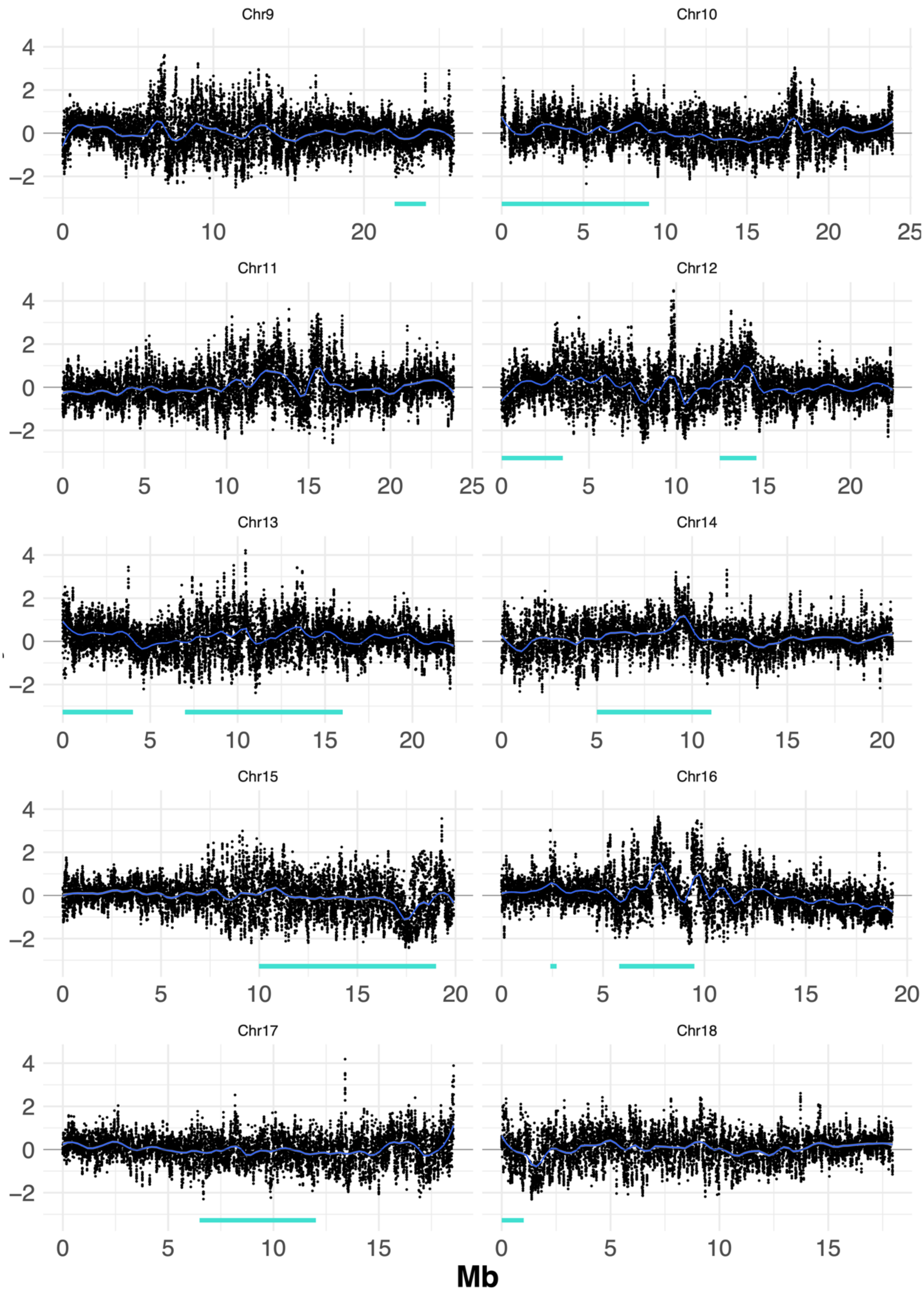

**Figure S8.** Estimates of Tajima's D[6] in 25 Kb windows along the chromosomes of polar cod. Position of chromosomal inversions marked with a green bar. Analysis was run per sample location, and for visualization shown together. See

Supplementary html: Tajima's D for detailed estimates for each sample location.

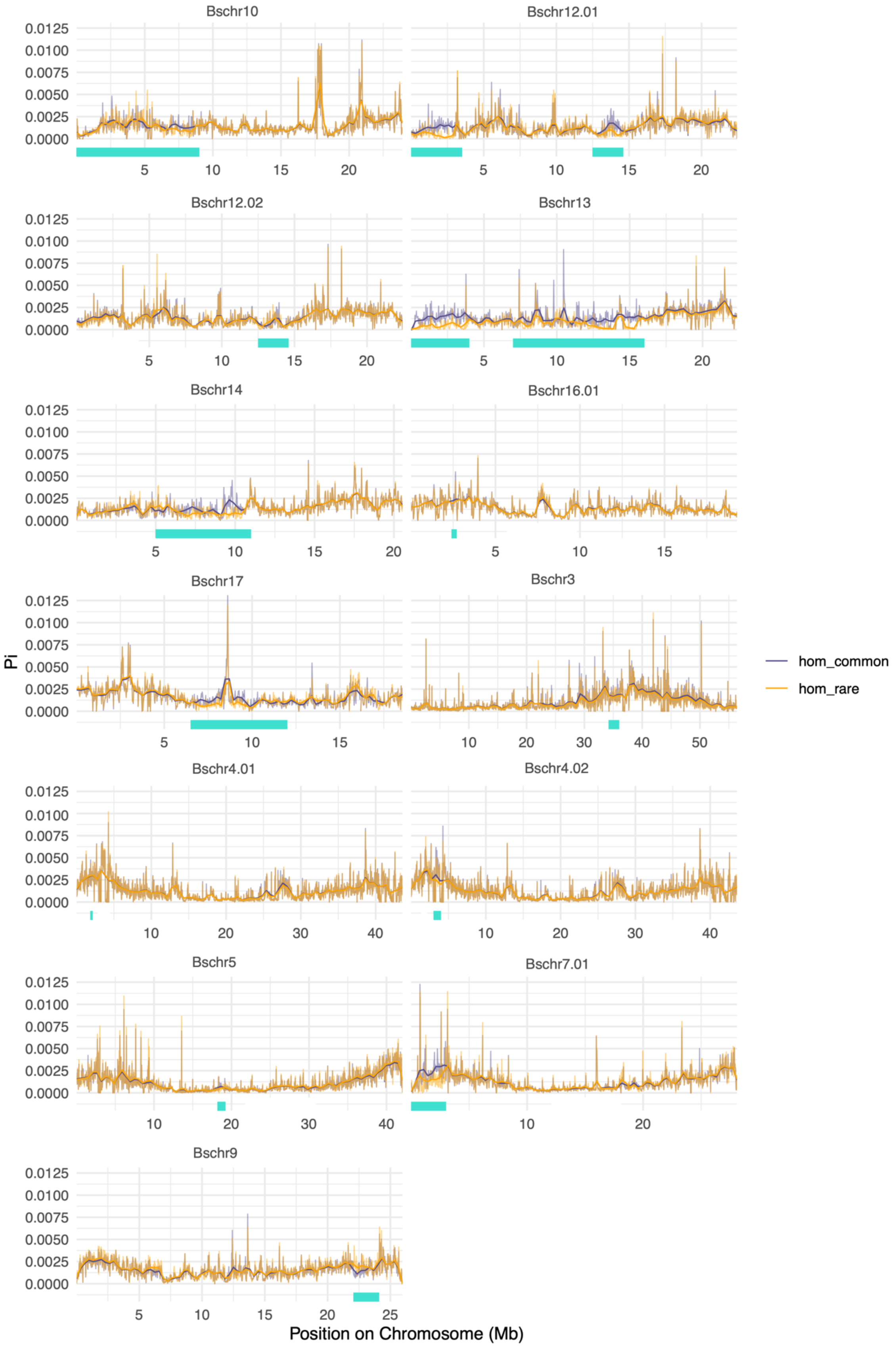

**Figure S9.** Estimates of Nucleotide diversity (π) by Pixy[7] in 25 kb windows along the chromosome, within each haplotype for a selection of inversions (which displayed three genotype clusters). The position of the inversions is marked with a green bar.

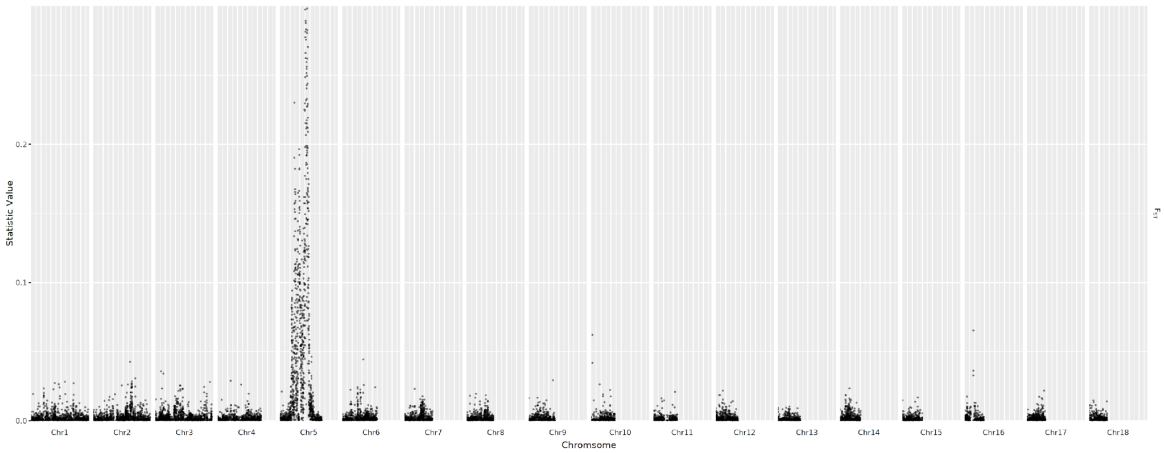

**Figure S10.** Genetic differentiation (estimated as $F_{ST}$) calculated in 25 kb widows using Pixy[7], between morphologically determined males and females across all chromosomes.

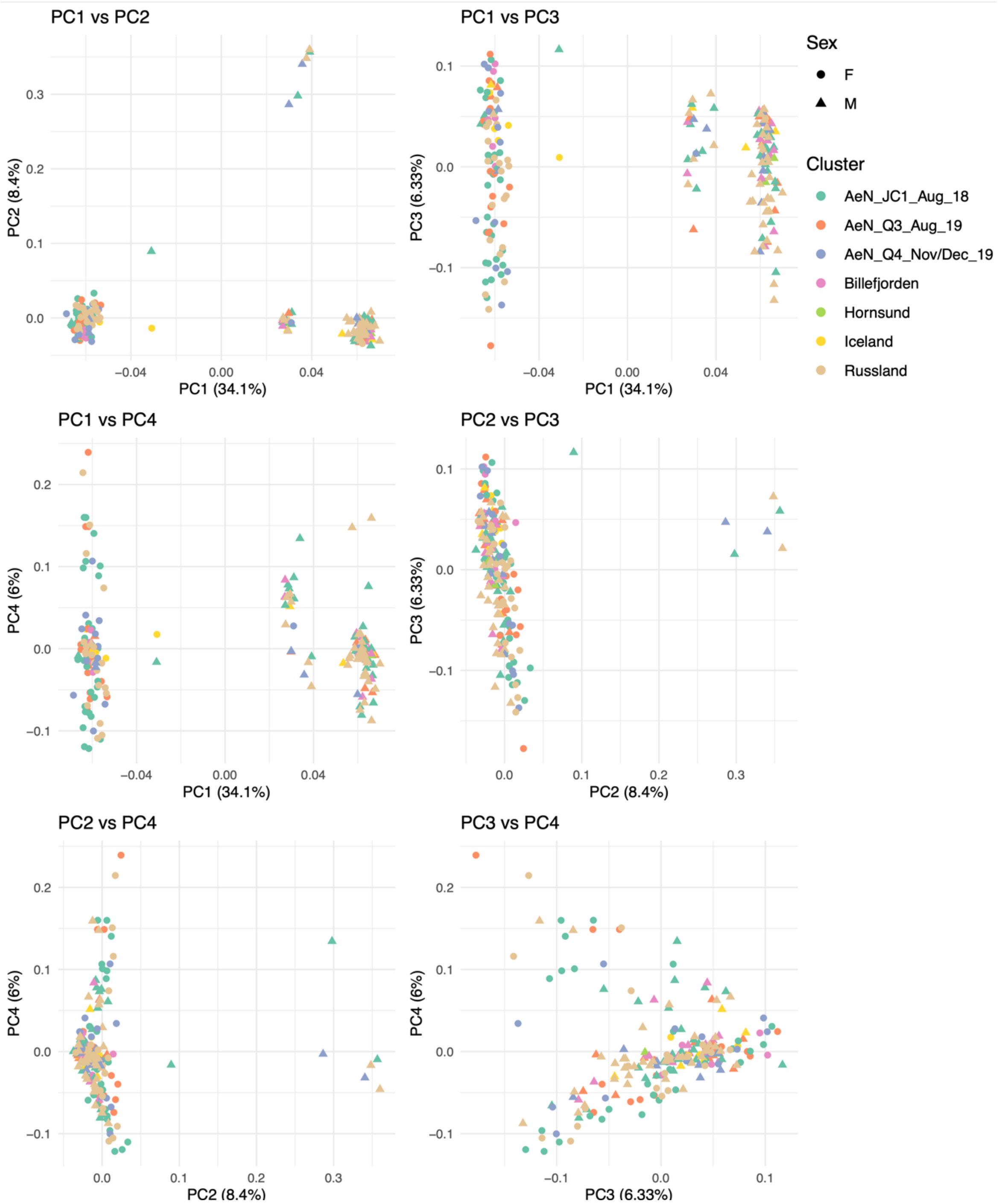

**Figure S11.** PCA of SNPs within the putative sex-determining region. Differentiation between sexes was found, as well as sub-structuring of male polar cod, making up a large and a small cluster along PC1. The male clusters do not seem to be associated with geographic location of samples.

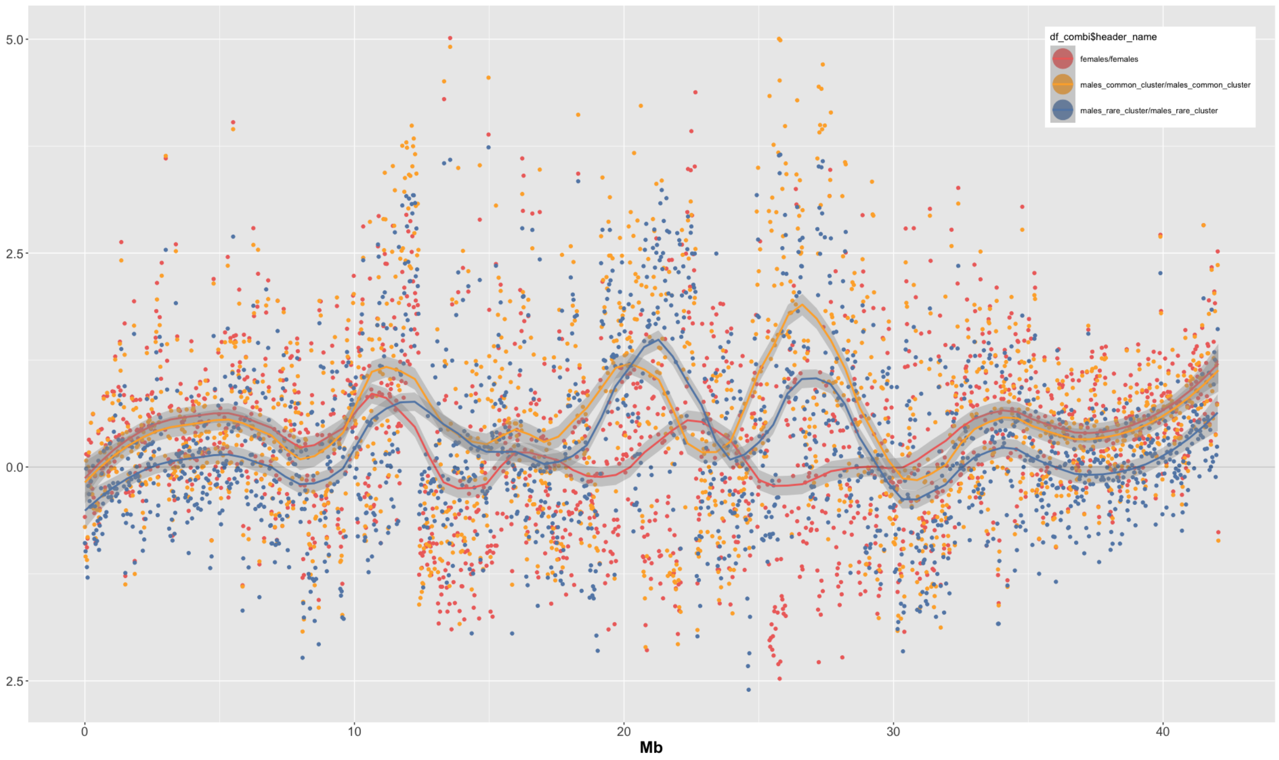

**Figure S12.** Estimates of Tajima's D in 25 kb windows along chromosome 5 performed for females, and the two male clusters, separately.

### Supplementary Materials and Methods

#### Specimen collection

For the Northwest Barents Sea region, samples were collected in a south-to-north gradient as part of Nansen Legacy research cruise transect (https://arvenetternansen.com/nansen-legacy-main-transect/). The initial sampling was conducted in August 2018, then subsequently repeated in August and December-November 2019. For all specimens sampled, gill and fin tissues were collected and preserved on 96% EtOH, and long-term stored at -80 °C for downstream genomic extraction and analyses. Along with the tissue samples, phenotypic measurements such as weight (g), length (mm), and sex (M or F) determined by visual inspection of gonads were recorded. Additionally, otoliths were collected for all specimens for the purpose of age determination conducted by the Institute of Marine Research, Bergen, Norway. The samples from the Northeast Barents Sea were collected in October 2019, from five sampling locations in a north-south gradient along the coast of Novaya Zemlya, including a northernmost location in the Kara Sea (Figure 1A, S8 and Table S9-17 for metadata sheets). We also included samples from three fjords on the western coast of Svalbard: Hornsund, Billefjorden, and Kongsfjorden, as well as specimens caught off the coast of Iceland in 2012. Finally, the dataset was supplemented with a few samples each from Tyrolerfjord, Greenland, and Davis Strait, Canada (see Table S8).

The specimens selected for sequencing were balanced to be close to equal males and females. Moreover, by estimating the age for most samples we found that the majority of our samples were one and two-year old (Figure S14). We found that the different sample locations also made up some differences in year classes represented: Iceland, Northwest Barents Sea summer 2018, and Northeast Barents Sea autumn 2019 consists of one-, two- and three-year-old fish, whilst other sample groups were found to contain generally younger fish. See Figure S14 for summary plots of age and sex distribution.

All samples used in this study have been collected in a responsible manner in connection to research surveys (as part of larger hauls for stock assessments). The fish were humanely sacrificed before sampling in accordance with the guidelines set by national and international animal welfare laws (e.g., www.norecopa.no), and thus no specific legislation was needed.

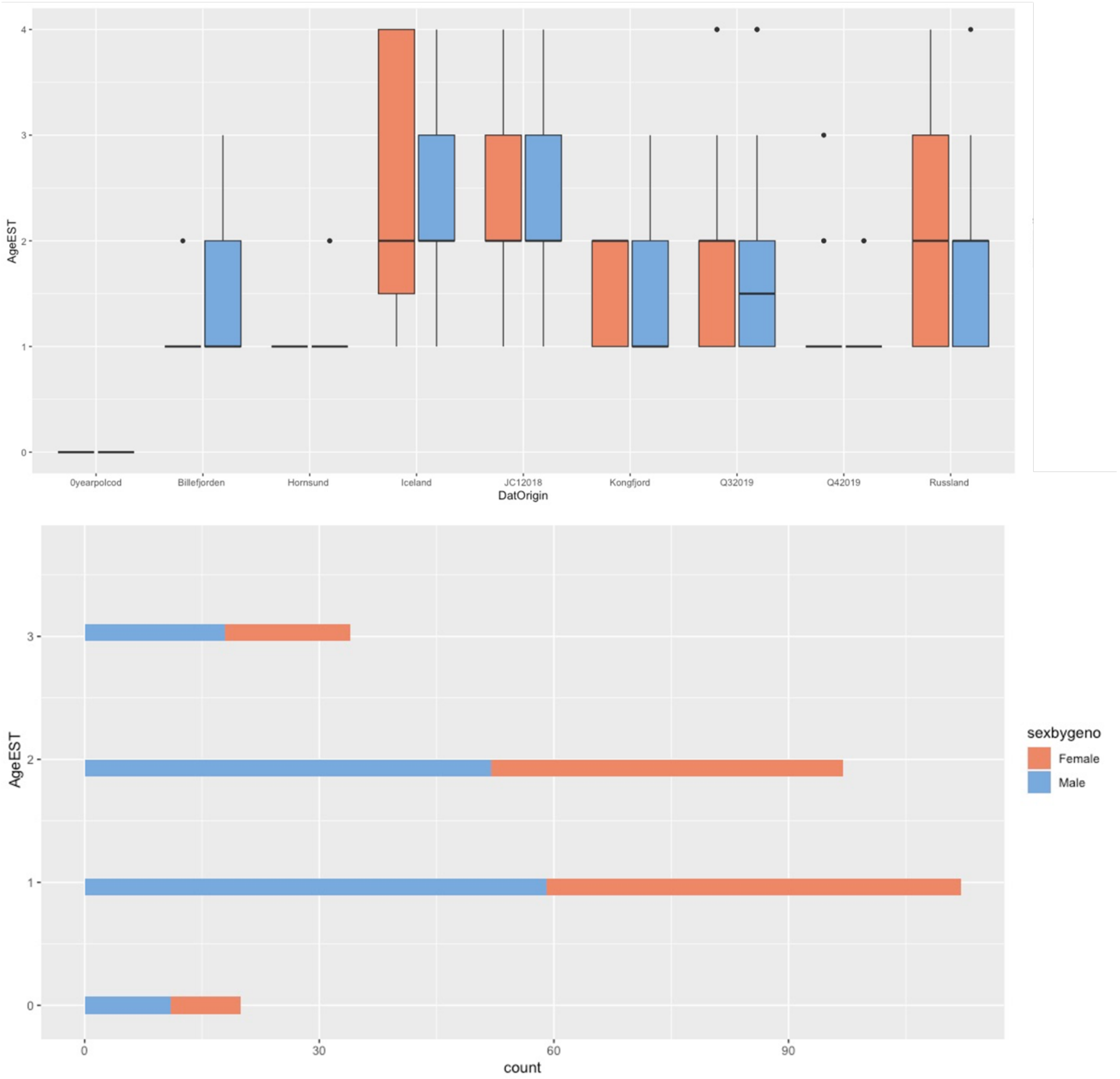

**Figure S13.** Summary of the age distribution of the specimen collection. JC1=Northwest Barents Sea summer 2018, Q3=Northwest Barents Sea summer 2019 and Q4=Northwest Barents Sea winter 2019. Among 258 polar cod (the majority of specimens collected JC1, Q3, and Q4), 127 (50 %) were aged using otolith reading. To estimate the age of the 127 remaining polar cod, we used available information on individual weight and length by comparing them to the 127 fish of known age, weight, and length. We used the following criteria: 1 year of age for fish <13 g and <13 cm long; 2 years of age for fish with weight >=13g and <30.3g with a length >=13 cm and <16 cm; 3 years of age for fish with weight >= 30.3 g and <45 g and length >=16cm and <19.3cm; 4 years of age for fish with weight >=45g and length > 19.3cm. 21 polar cod were not aged using this method. Among the 24 polar cod not aged using this method, 6 were aged visually.

**DNA quantification, library preparation, and genome sequencing**

The population sequencing was performed in three rounds (in 2018, 2019, and 2021) of sequencing at the same facility (Norwegian Sequencing Centre, Oslo). The sequencing was done using the same methods across all samples, with minor differences only in PCR cycles (noted below) as well as an extra round of sequencing performed for the 2018 samples that yielded slightly higher mapping depth for these samples (see Figure S14B).

The DNA samples were quantified using the FLUOStar Optima (BMG Labtech) with the Qubit dsDNA HS Assay Kit chemistry (ThermoFisher Scientific). Normalization of all DNA samples to 20ng/ul with Elution Buffer (Qiagen) was performed using the Sciclone G3 NGS Workstation (Perkin Elmer). Normalized DNA samples were sheared using the E220 focused-ultrasonicator (Covaris) with the appropriate manufacturer´s settings for a target fragment mean size of 350 bp. After shearing all samples were purified and size selected using KAPA Pure beads (Roche) in a ratio 0.8x (beads:sample) in order to remove fragments shorter than 200 bp prior to library preparation. Library preparation was performed using the KAPA Hyper kit (Roche) on Mosquito LV (Low Volume) pipetting robot (sptlabtech). The library preparation reactions (End repair, A-tailing, and adapter Ligation) were performed using 5x reduced volume compared to the kit reaction volumes. The IDT for Illumina TruSeq DNA UD 96 Indexes (Illumina) were used for barcoding each 96 plate of samples. After the ligation of adapters, the samples volume was increased to 25ul with the addition of EB buffer (Qiagen), and one round of bead cleanup with ratio 0.8x was performed. The libraries were subsequently amplified with either 4 (samples sequenced in 2020 and 2021) or 5 (samples sequenced in 2018) cycles of PCR. The PCR reactions were done in 2x reduced volume compared to the kit PCR reactions. All incubations were executed according to manufacturer´s instructions. The final libraries were purified, and size selected using KAPA Pure beads (Roche) in a ratio 0.8x. After library preparation and cleanup, all libraries were run on a 5200 Fragment Analyzer System (Agilent) using the NGS Fragment Kit: DNF-473-0500 (Agilent) for determination of the average size of each library. Subsequently, absolute quantification of each library was done using the KAPA Library Quanitification Kits (Roche), on a LightCycler 480 qPCR instrument (Roche) in 10ul reaction volume. Finally, after determining the absolute concentration of each library (in nM) using the Fragment analyzer and qPCR results, all libraries were normalized to the same molarity using the Sciclone G3 NGS Workstation (Perkin Elmer) and equal volumes of each sample were pooled, creating 96plex pools. Each of the pools was sequenced on several lanes of a HiSeq4000 System (Illumina) in 2x150bp mode (150bp Paired End), using a HiSeq 3000/4000 SBS Kit (300 cycles) (Illumina).

**Mapping of sequence data, variant calling, and quality assessment**

Variant calling was performed for all samples combined (a total of 290 polar cod). The raw Illumina PE reads were trimmed for low-quality bases using Trimmomatic v0.39[8] with the following options and settings: ILLUMINACLIP:${ADAPT_FAST}:2:30:10, LEADING:5, TRAILING:5 SLIDINGWINDOW:5:10 ,MINLEN:50.

Trimmed reads were aligned to the *Boreogadus saida* reference genome (borSai1.0)[1] using the Burrows-Wheeler Alignment Tool (BWA) mem v0.7.17[9]. Alignment files were sorted and merged using SAMtools v1.9[10], and Picard MarkDuplicates v2.22.1[11] was used to identify and mark duplicated reads. Variant discovery, calculations of variant statistics, and hard filtering of variant sites were performed using the Genome Analysis Toolkit (GATK) v4.2.0[12]. First, variants were called within each individual separately using HaplotypeCaller, generating per individual GVCFs. GVCF's for each of the 290 samples were then imported into a GenomicsDataBase using the GenomicsDBImport tool. Joint genotyping was performed using the GenotypeGVCFs tool to produce final population-level genotype VCF's for each chromosome.

Raw variants were first split using GATK SelectVariants with settings -select-type SNP and -select-type NO_VARIATION resulting in variant files containing only SNPs and invariant sites. SNP variants were then manually quality assessed using the GATK v4.2.0[12] program "VariantsToTable" and VCFtools v0.1.16[13], visualization was done in R v4.0.3[14].

While initially filtering the whole genome SNP set, we uncovered a pattern of potential batch-related bias in the dataset (when the dataset was filtered and subsetted to neutral variants — not detected otherwise). By manual inspection of QualByDepth (QD) outputted from GATK v4.2.0 "VariantsToTable" we detected an excess of heterozygous sites (with low-quality scores) (Figure S16). We suspect this skewness in variants — which was found to contribute to a bias pattern in our dataset, likely stems from batch 1 being applied one extra round of qPCR cycle, compared to batch 2 and batch 3. We therefore adjusted the QD filter setting to QD < 5.0 and found that the bias was corrected (Figures S17 and S18).

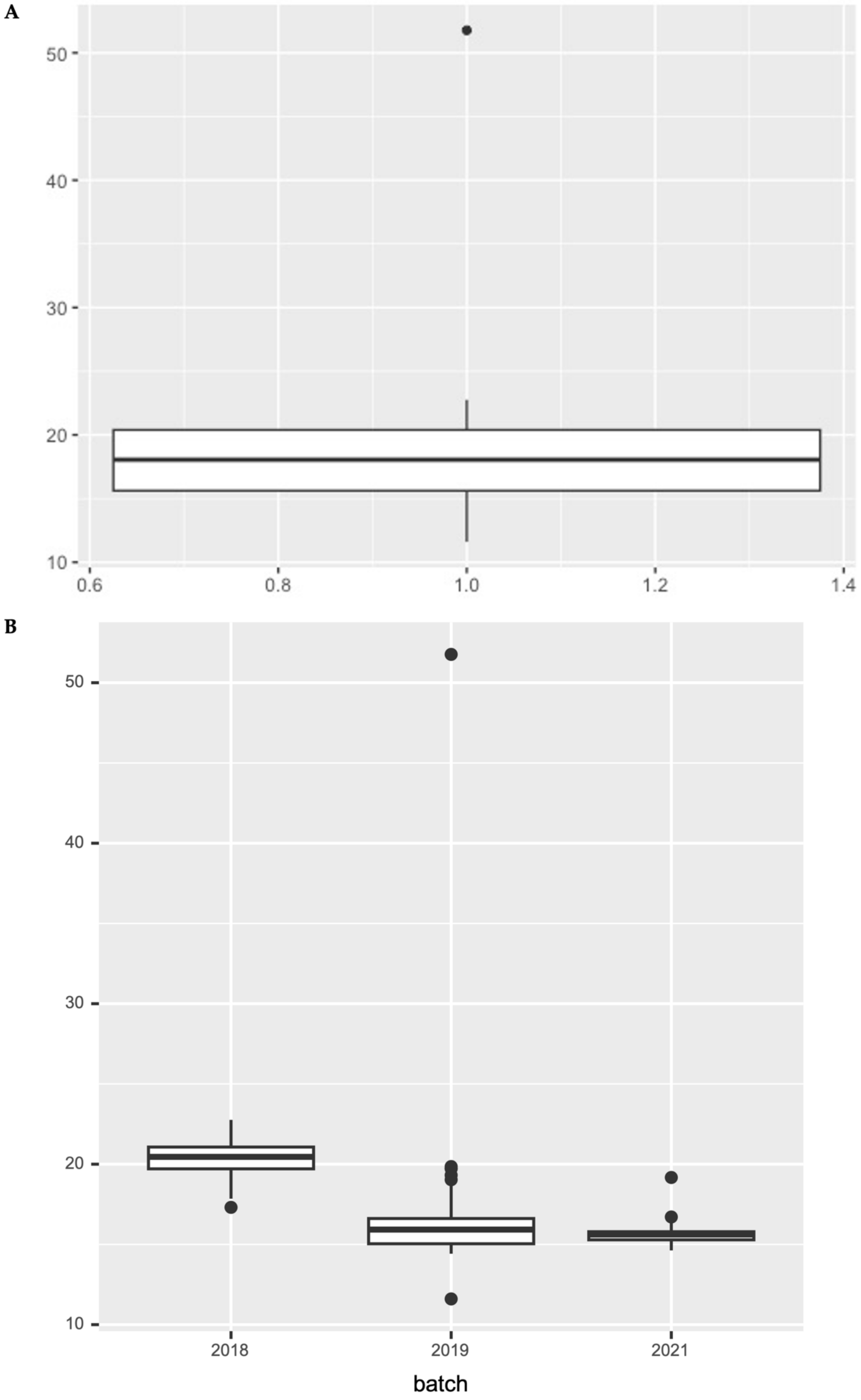

**Figure S14. A)** Individual read mapping depth across 290 samples. **B)** Individual read mapping depth where samples are divided by batch of sequencing. The first batch (2018) was sequenced slightly deeper than the subsequent batches. The middle line (in the boxes) denotes the median value. Box indicates interquartile range.

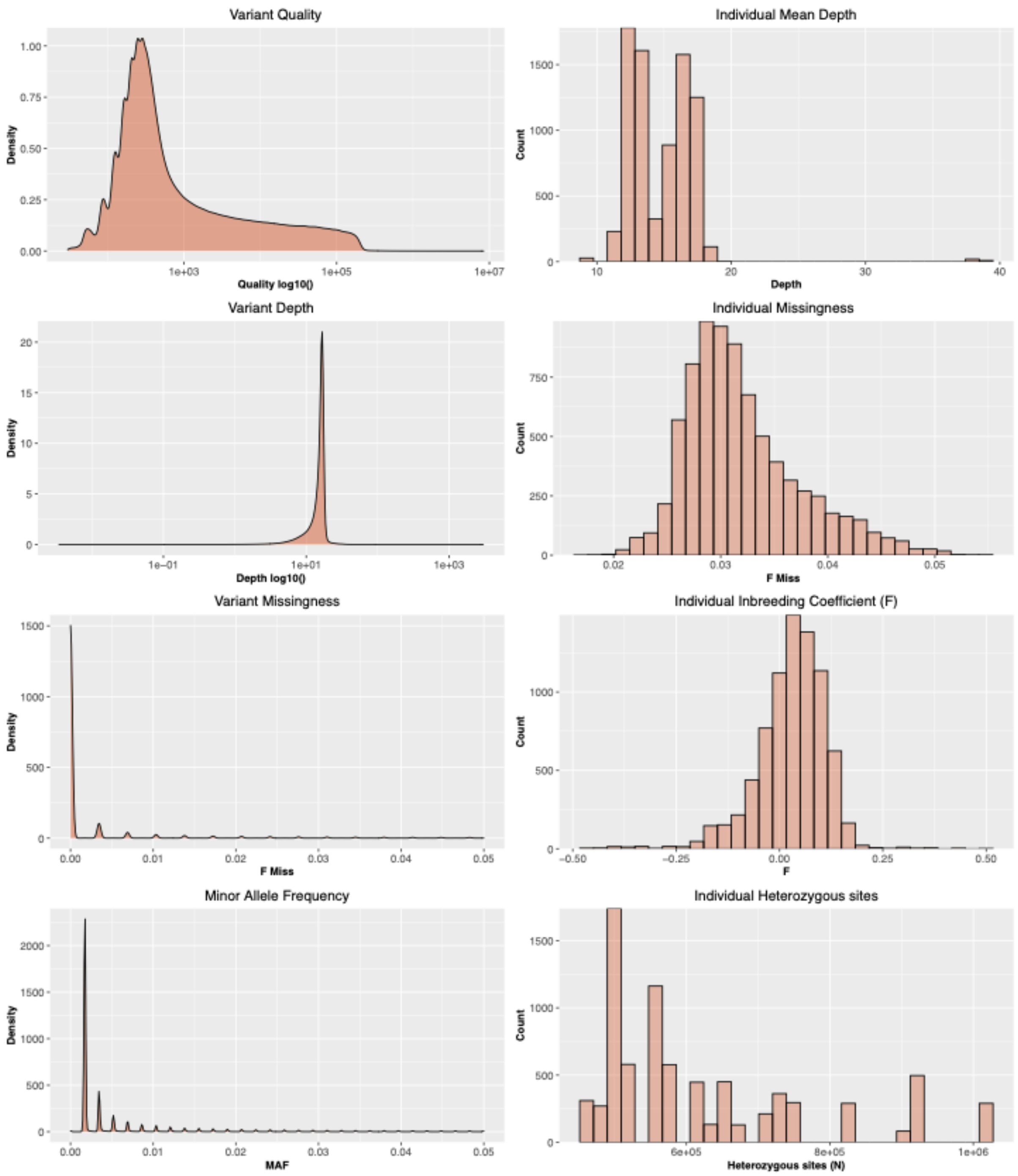

**Figure S15.** Variant quality before filtering summarized using VCFtools[13].

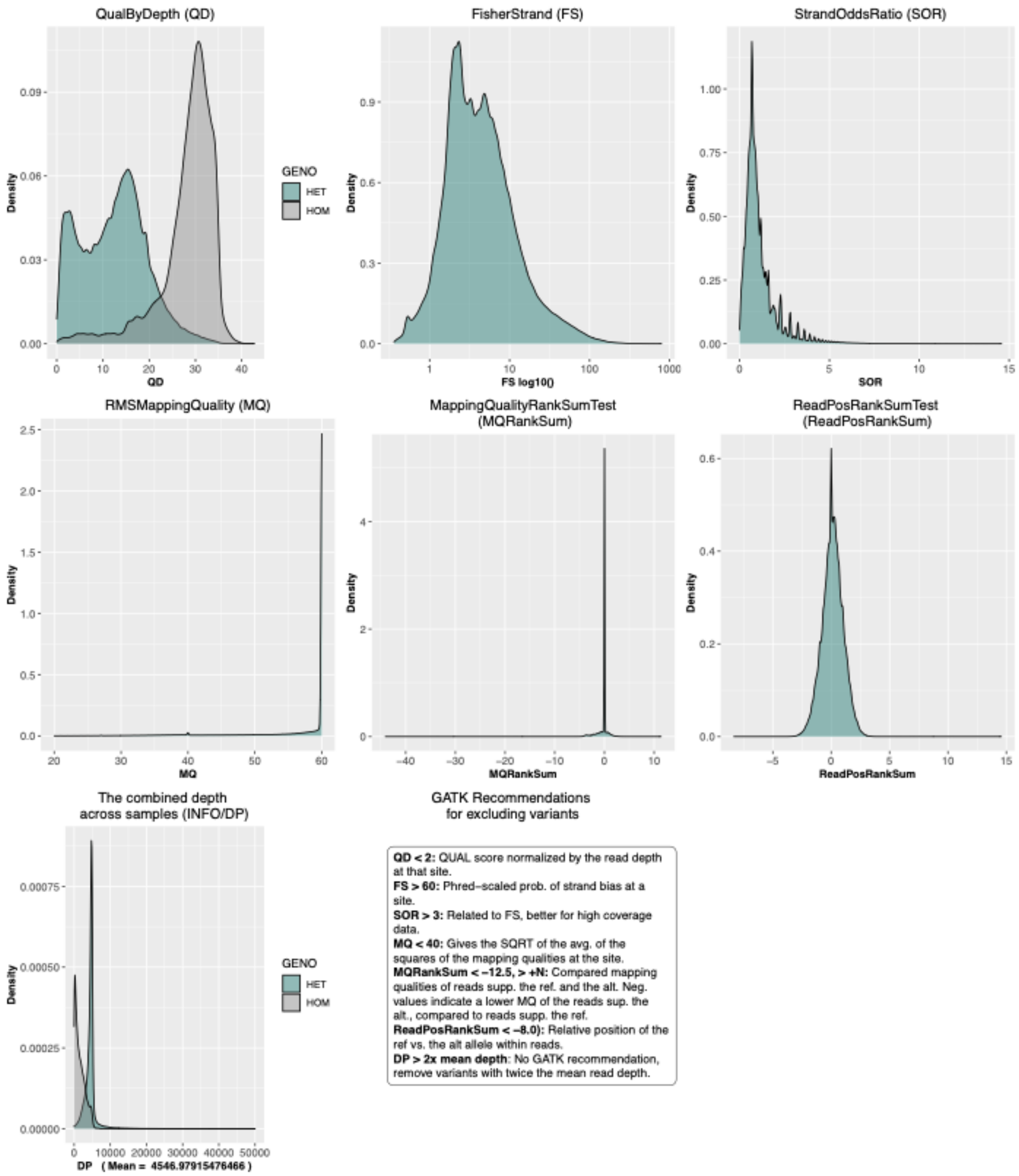

**Figure S16.** Variant quality before filtering summarized by GATK tool "variant-to-table", revealed an elevated number of het sites with low-quality scores (QD plot).

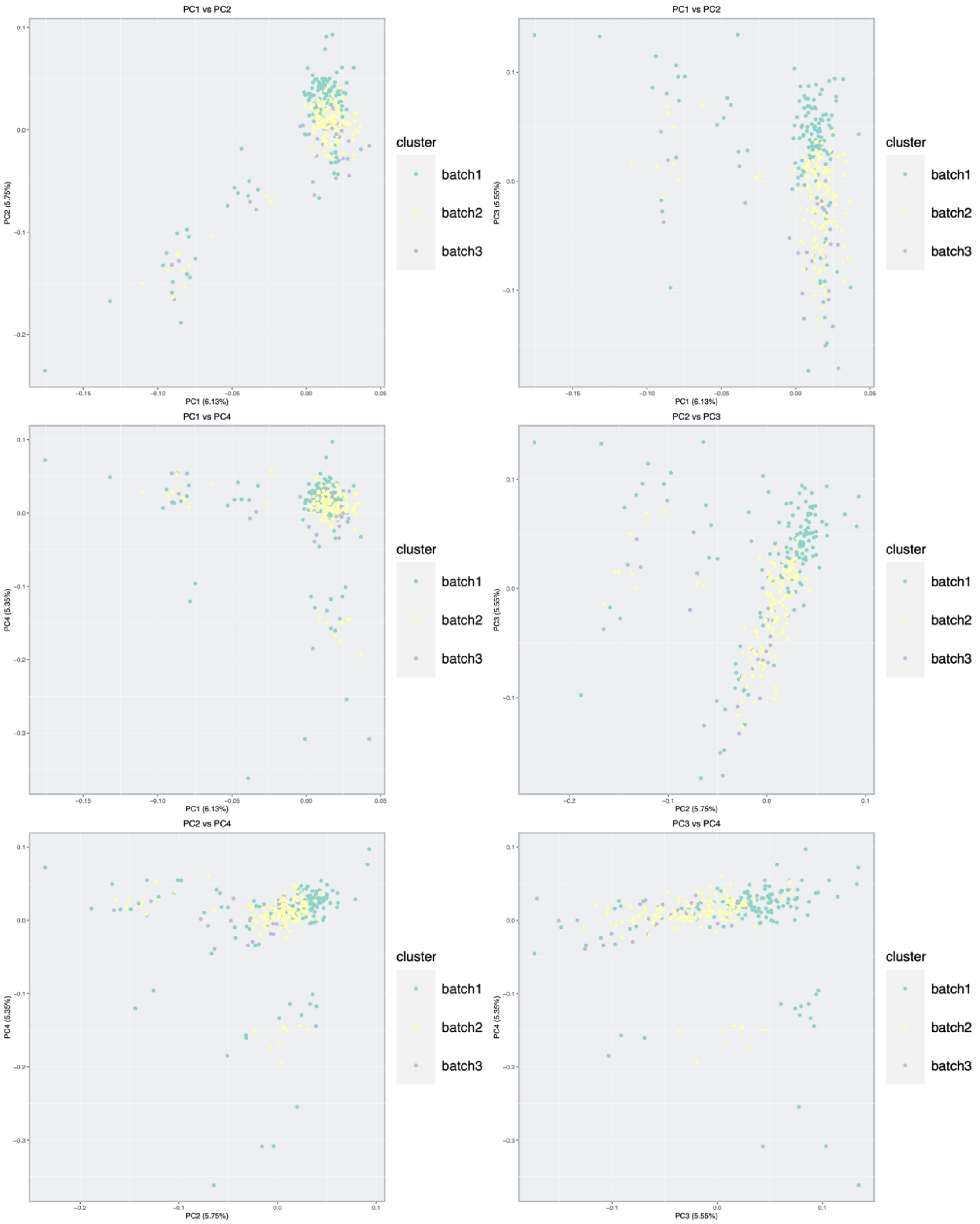

**Figure S17.** PCA across A genome-wide SNP set during filtering evaluation, where large, linked regions were removed, and a potential batch-related bias became apparent.

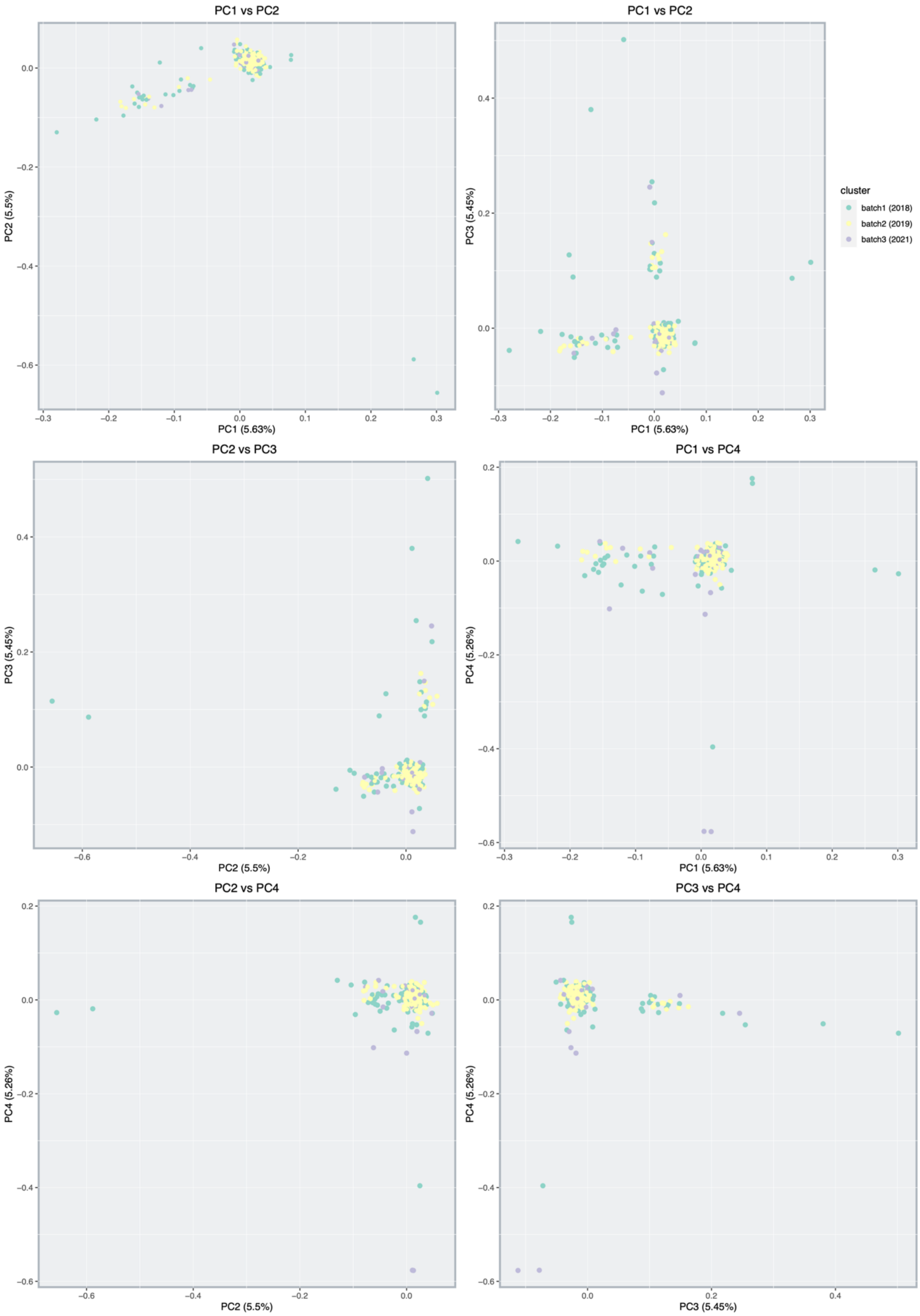

**Figure S18.** PCA across the final putative neutral SNP dataset after additional quality filtering, indicates the bias was successfully removed.

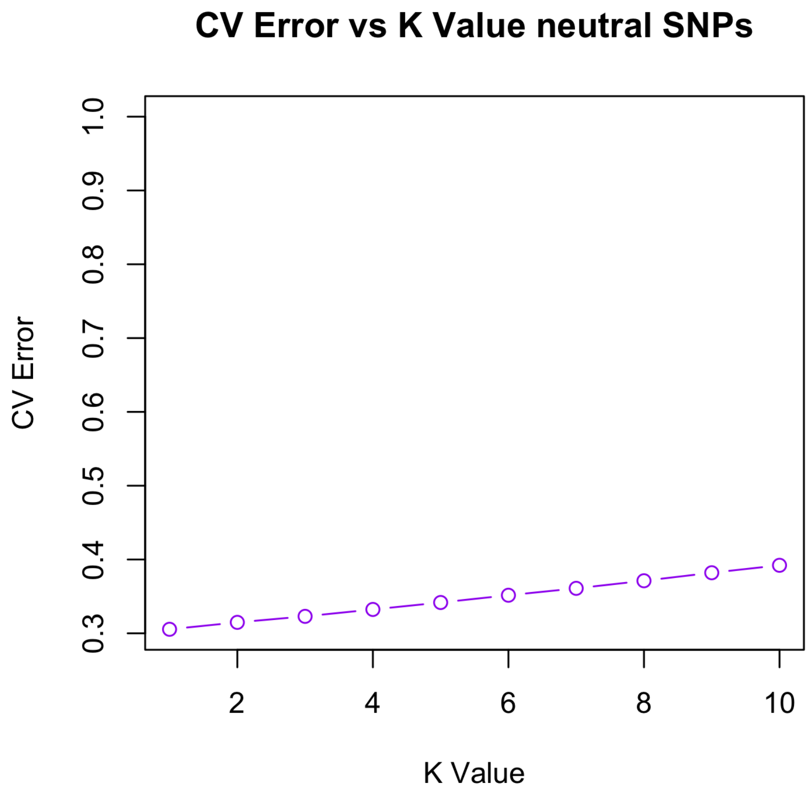

**Figure S19.** Graph showing CV error estimates across ADMIXTURE[15] analyses K=1 through 10 using the 564 970 neutral SNPs as input data. The lowest estimate was observed for K=1, followed by a gradual increase in estimate.

**Estimated female effective population size and mitochondrial phylogeny among a selection of the samples**

Reads mapping to the mitogenomes in proper pairs against polar cod and Arctic cod for selected samples (N=92 from the Barents Sea) were extracted and sorted by read name using SAMtools v1.9[10]. Bam files were converted back to FASTQ format using BEDTools v2.27.1[16] bamtofastq tool. Assembly of mitogenomes was performed using MitoFinder v1.4.1[17] with default settings and the assembler MEGAHIT v1.0[18,19]. MiTFi v0.1[20] was used to annotate the mitochondrial tRNAs. Mitogenome annotation was performed using the available mitogenome references for Arctic cod (NCBI RefSeq: NC_010122.1) and polar cod (NCBI RefSeq: NC_010121.1) as MitoFinder requires annotated mitogenomes in GenBank reference format.

A maximum likelihood (ML) phylogeny was inferred to assess if the samples used for demographic inference formed good clades, e.g., assigned to the correct species and not showing strong population structure. For the ML phylogeny, we included the polar cod samples, the Atlantic cod NEAC and NCC mitogenomes.

Atlantic haddock (RefSeq: NC_007396.1), and burbot[1] were used as outgroups. PCGs, excluding *COX2* as this was missing for one sample, were concatenated using PhyKIT v1.11.7[1] and aligned with MAFFT v7.453[21]. ML inference was run in IQ-Tree2 v2.2.0[22], and the substitution model was inferred using MFP[23]. Bootstrap support was estimated with 1,000 replicates of UFBoot[24]. The inference resulted in a phylogenetic topology delineating three main clades for polar cod (Figure S21).

Of the 92 samples of polar cod, only one failed assembly due to MitoFinder assembling two potential mitogenomes. Also, not all samples had complete PCGs. Thus, these samples were removed leaving 84 samples for demographic inference with all 13 complete PCGs. Annotated PCG from each individually MitoFinder assembled intraspecific level mitogenome of polar cod were extracted and aligned using MAFFT v7.453 were concatenated with PhyKIT v1.11.7 create_concat to produce a supermatrix.

The female effective population size ($N_e$) was estimated in BEAST v2.6.7[25] under the Bayesian skyline model[26]. The substitution model was in-ferred using bModelTest with the namedExtended list of models. The coalescent Bayesian skyline prior was applied under a strict clock with a rate of 1.14x10-8 substi-tution/site/year used, as reported for Atlantic cod[27]. This rate was applied because there are no estimates for the substitution rate of polar cod. The analysis was performed using a chain length of 800,000,000, and sampling was done every 1,000 itera-tions with bPopSizes and bGroupSizes set to 5 dimensions. Tracer v1.7.2[28] was used to check convergence and to reconstruct the Bayesian skyline with default settings. (sky-line variant = stepwise (constant), maximum time to root height = lower 95% HPD, root height = Treeheight, number of bins = 100).

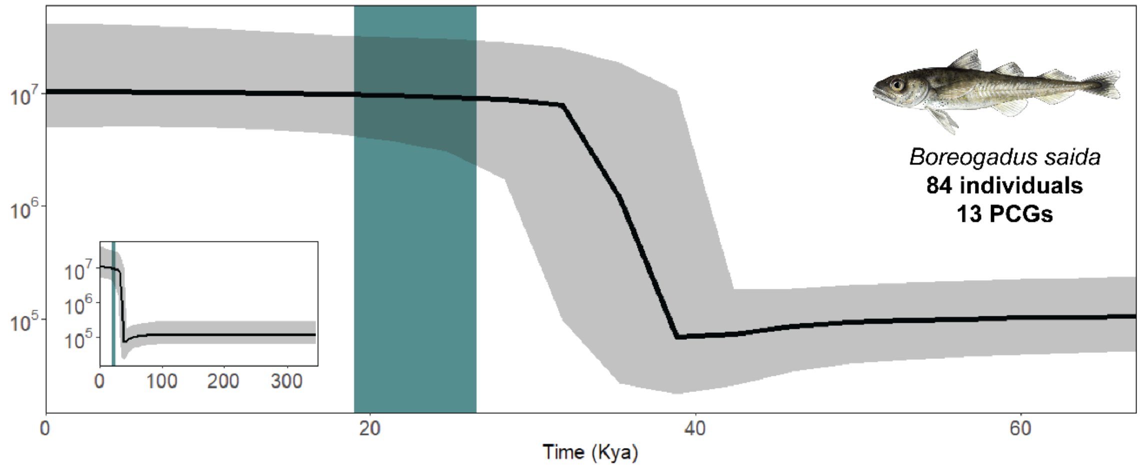

**Figure S20.** Inferred female effective population size (Ne) for polar cod using all 13 PCGs and 84 specimens with complete genes. The smaller panel represents the total inferred coalescence time for polar cod. The blue bar indicates the last glacial maximum. The mitochondrial demographic inference for polar cod using 84 individuals revealed a slight decrease in female Ne around 40 Kya followed by a steep increase until 30 Kya. The median estimated Ne at year 0 (present time) was around ~$10^7$.

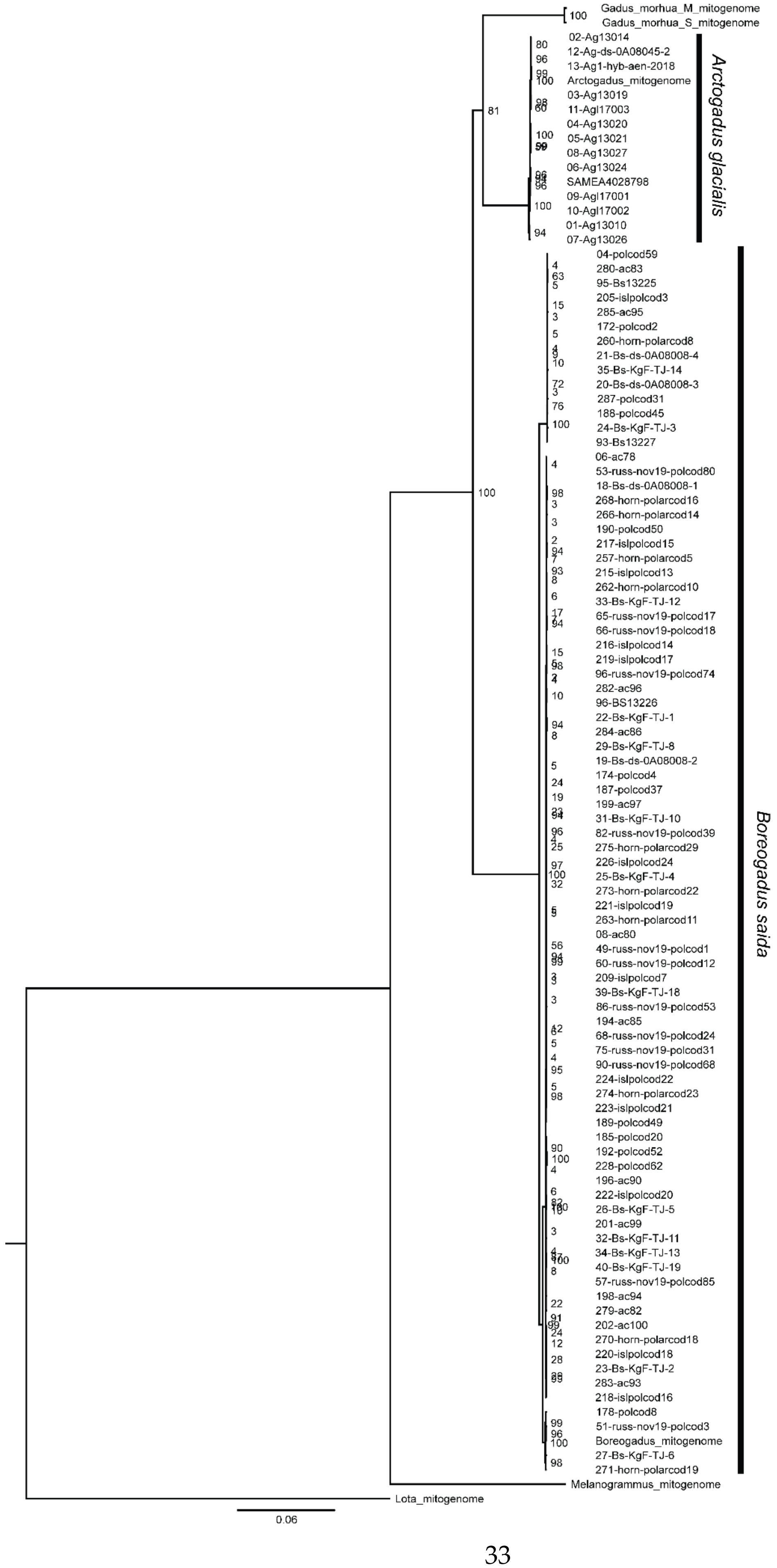

Gadus_morhua_M_mitogenome
Gadus_morhua_S_mitogenome
02-Ag13014
12-Ag-ds-0A08045-2
13-Ag1-hyb-aen-2018
Arctogadus_mitogenome
03-Ag13019
11-Agl17003
04-Ag13020
05-Ag13021
08-Ag13027
06-Ag13024
SAMEA4028798
09-Agl17001
10-Agl17002
01-Ag13010
07-Ag13026
Arctogadus glacialis
04-polcod59
280-ac83
95-Bs13225
205-islpolcod3
285-ac95
172-polcod2
260-horn-polarcod8
21-Bs-ds-0A08008-4
35-Bs-KgF-TJ-14
20-Bs-ds-0A08008-3
287-polcod31
188-polcod45
24-Bs-KgF-TJ-3
93-Bs13227
06-ac78
53-russ-nov19-polcod80
18-Bs-ds-0A08008-1
268-horn-polarcod16
266-horn-polarcod14
190-polcod50
217-islpolcod15
257-horn-polarcod5
215-islpolcod13
262-horn-polarcod10
33-Bs-KgF-TJ-12
65-russ-nov19-polcod17
66-russ-nov19-polcod18
216-islpolcod14
219-islpolcod17
96-russ-nov19-polcod74
282-ac96
96-BS13226
22-Bs-KgF-TJ-1
284-ac86
29-Bs-KgF-TJ-8
19-Bs-ds-0A08008-2
174-polcod4
187-polcod37
199-ac97
31-Bs-KgF-TJ-10
82-russ-nov19-polcod39
275-horn-polarcod29
226-islpolcod24
25-Bs-KgF-TJ-4
273-horn-polarcod22
221-islpolcod19
263-horn-polarcod11
08-ac80
49-russ-nov19-polcod1
60-russ-nov19-polcod12
209-islpolcod7
39-Bs-KgF-TJ-18
86-russ-nov19-polcod53
194-ac85
68-russ-nov19-polcod24
75-russ-nov19-polcod31
90-russ-nov19-polcod68
224-islpolcod22
274-horn-polarcod23
223-islpolcod21
189-polcod49
185-polcod20
192-polcod52
228-polcod62
196-ac90
222-islpolcod20
26-Bs-KgF-TJ-5
201-ac99
32-Bs-KgF-TJ-11
34-Bs-KgF-TJ-13
40-Bs-KgF-TJ-19
57-russ-nov19-polcod85
198-ac94
279-ac82
202-ac100
270-horn-polarcod18
220-islpolcod18
23-Bs-KgF-TJ-2
283-ac93
218-islpolcod16
178-polcod8
51-russ-nov19-polcod3
Boreogadus_mitogenome
27-Bs-KgF-TJ-6
271-horn-polarcod19
Boreogadus saida
Melanogrammus_mitogenome
Lota_mitogenome
0.06

**Figure S21.** Mitochondrial phylogeny among a selection of polar cod samples from the Barents Sea, revealing three main clades. Additionally, 15 Arctic cod samples as well as burbot and Atlantic haddock set as outgroups.

#### Supplementary Tables

**Table S1.** Pairwise global $F_{ST}$ estimates from comparisons between Northwest Barents Sea summer 2018, 0-group fish, Billefjorden, Northeast Barents Sea autumn 2019, as well as two individual Northeast Barents Sea locations, vs all other sampling locations. Estimates were calculated as a global mean from measurements using Pixy[7] in 25 kb windows. See Supplementary html: $F_{ST}$ for window-based results across chromosomes.

| | NWBS 2018 | zerogroup | Billefjorden | NEBSA 2019 | St18 | St9 |
|---|---|---|---|---|---|---|
| **Davis_strait** | 0,01633 | 0,01432 | 0,01558 | 0,01599 | 0,01501 | 0,01612 |
| **Greenland** | 0,00128 | 0,00000 | 0,00148 | 0,00092 | 0,00062 | 0,00147 |
| **Iceland** | 0,00117 | 0,00000 | 0,00088 | 0,00131 | 0,00106 | 0,00080 |
| **NWBS 2018** | | 0,00000 | 0,00043 | 0,00063 | 0,00195 | 0,00128 |
| **NWBSs 2019** | 0,00073 | 0,00000 | 0,00079 | 0,00013 | 0,00006 | 0,00020 |
| **NWBSw 2019** | 0,00000 | 0,00000 | 0,00016 | 0,00000 | 0,00000 | 0,00024 |
| **zerogroup** | 0,00000 | | 0,00000 | 0,00000 | 0,00021 | 0,00010 |
| **Billefjorden** | 0,00043 | 0,00000 | | 0,00031 | 0,00016 | 0,00023 |
| **Kongsfjorden** | 0,00109 | 0,00000 | 0,00010 | 0,00043 | 0,00000 | 0,00032 |
| **Hornsund** | 0,00058 | 0,00000 | 0,00036 | 0,00000 | 0,00029 | 0,00000 |
| **St40** | | 0,00058 | | | 0,00107 | 0,00175 |
| **St22** | | 0,00000 | | | 0,00000 | 0,00112 |
| **St18** | | 0,00021 | | | | 0,00000 |
| **St17** | | 0,00000 | | | 0,00000 | 0,00110 |

NWBS 2018=Northwest Barents Sea summer 2018
NWBSs 2019=Northwest Barents Sea summer 2019
NWBSw 2019=Northwest Barents Sea winter 2019
NEBSA 2019=Northeast Barents Sea autumn 2019 (all stations together)
St=Sampling stations in the Northeast Barents Sea autumn 2019 (see Figure 2C for locations of stations).

**Table S2.** Overview of genomic structures identified in the genome of polar cod. *=Inversions identified that display three genotype statuses (i.e., Hom common, Het, Hom rare). **=Inversions found to display three discrete homozygous genotypes.

| Chromosome | Inversion name | Start position (bp) | End position (bp) | Size (bp) | Annotated genes in the region (n) | Structure type |
|---|---|---|---|---|---|---|
| Chr2 | Bschr2 | 23000000 | 38000000 | 15000000 | 783 | Complex Inversion* |
| Chr3 | Bschr3 | 34200000 | 36000000 | 1800000 | 122 | Classical inversion* |
| Chr4 | Bschr4.01 | 1840000 | 2150000 | 310000 | 29 | Classical inversion* |
| Chr4 | Bschr4.02 | 3000000 | 4000000 | 1000000 | 71 | Classical inversion* |
| Chr5 | Bschr5 | 18200000 | 19200000 | 1000000 | 86 | Classical inversion* |
| Chr5 | | 22000000 | 30000000 | 8000000 | | Sex-determining haploblock |
| Chr6 | Bschr6.01 | 19000000 | 23500000 | 4500000 | 220 | Complex Inversion** |
| Chr6 | Bschr6.02 | 23000000 | 25500000 | 2500000 | 114 | Complex Inversion** |
| Chr7 | Bschr7.01 | 1 | 3000000 | 2999999 | 275 | Complex inversion* |
| Chr7 | Bschr7.02 | 13000000 | 17500000 | 4500000 | 229 | Complex Inversion** |
| Chr9 | Bschr9 | 22050000 | 24100000 | 2050000 | 154 | Classical inversion* |
| Chr10 | Bschr10 | 1 | 9000000 | 8999999 | 769 | Classical inversion* |
| Chr12 | Bschr12 | 1 | 3500000 | 3499999 | 220 | Classical inversion* |
| Chr12 | Bschr12 | 12500000 | 14600000 | 2100000 | 169 | Complex inversion* |
| Chr13 | Bschr13 | 1 | 4000000 | 3999999 | 670 | Classical inversion (part* A) |
| Chr13 | | 7000000 | 16000000 | 9000000 | 304 | Classical inversion (part B) |
| Chr14 | Bschr14 | 5000000 | 11000000 | 6000000 | 447 | Classical inversion* |
| Chr15 | Bschr15 | 10000000 | 19000000 | 9000000 | 611 | Complex Inversion** |
| Chr16 | Bschr16.01 | 2400000 | 2700000 | 300000 | 33 | Classical inversion* |
| Chr16 | Bschr16.02 | 5800000 | 9500000 | 3700000 | 315 | Complex Inversion** |
| Chr17 | Bschr17 | 6500000 | 12000000 | 5500000 | 354 | Classical inversion* |
| Chr18 | Bschr18 | 1 | 1000000 | 999999 | 74 | Complex inversion* |

**Table S3.** Results for correlation analysis using Pearson's product-moment correlation between the different inversions displaying three genotype clusters, where we tested for associations between pairs of inversions.
Table provided online, as an Excel sheet:
https://www.mn.uio.no/ibv/personer/vit/snhoff/table_s3_inversion_correlation_analysis.xlsx

**Table S4.** Tests for inversion genotypes and Hardy Weinberg equilibrium, and estimates of $F_{IS}$ for each listed inversion among all 290 specimens.

| Inversion | Chi2 | Df | PrChi2 | PrExact | $F_{IS}$ |
|---|---|---|---|---|---|
| Bchr10 | 2,51 | 1 | 0,113 | 0,101 | 0,094 |
| Bchr12.01 | 5,04 | 1 | 0,025 | 0,038 | 0,134 |
| Bchr13 | 3,10 | 1 | 0,078 | 0,12 | -0,105 |
| Bchr14 | 4,62 | 1 | 0,032 | 0,04 | -0,128 |
| Bchr16.01 | 0,71 | 1 | 0,398 | 0,47 | -0,050 |
| Bchr17 | 1,57 | 1 | 0,211 | 0,248 | 0,075 |
| Bchr3 | 5,37 | 1 | 0,021 | 0,021 | 0,138 |
| Bchr4.01 | 0,03 | 1 | 0,867 | 0,878 | 0,010 |
| Bchr4.02 | 5,37 | 1 | 0,020 | 0,025 | -0,138 |
| Bchr5 | 4,60 | 1 | 0,032 | 0,035 | 0,128 |
| Bchr7 | 0,57 | 1 | 0,449 | 0,65 | -0,045 |
| Bchr9 | 2,60 | 1 | 0,107 | 0,166 | -0,097 |
| Bchr2 | 0,61 | 1 | 0,436 | 0,479 | -0,046 |
| Bchr12.02 | 0,07 | 1 | 0,787 | 0,891 | -0,016 |
| Bchr18 | 0,62 | 1 | 0,432 | 0,544 | -0,047 |

**Table S5.** Hemoglobin and flanking gene searches and hits using Blast+[29] in the genome of polar cod.

| MN cluster / Query gene short name | Chromosome hit | Start position of hit | End position of hit | Strand | Hit identity score | note |
|---|---|---|---|---|---|---|
| ***POLR3K*** | Chr12 | 14646657 | 14647401 | + | 98-100 | |
| ***Mgrn1*** | Chr12 | 14655270 | 14661366 | + | 98-100 | |
| ***MPG*** | Chr12 | 14709928 | 14712007 | + | 92-98 | |
| ***nprl3*** | Chr12 | 14722709 | 14712829 | - | 98-100 | |
| ***Hbb5*** | Chr12 | 14726625 | 14727274 | + | 97< | |
| ***Hba1*** | Chr12 | 14728813 | 14728153 | - | 97< | |
| ***Hbb1*** | Chr12 | 14730427 | 14731485 | + | 97< | interval fits perfect with the annotation (gff3: HBB1) |
| ***Hba4*** | Chr12 | 14742272 | 14742942 | + | 97< | |
| ***kank2*** | chr12 | 14752237 | 14744959 | - | 97-100 | |
| **LA cluster/ Query gene short name** | **Chromosome hit** | **Start position of hit** | **End position of hit** | **strand** | **identities** | **note** |
| ***RHBDF1*** | | 14206026 | 14211965 | + | 99-100 | |

| | | | | | | |
|---|---|---|---|---|---|---|
| ***Hb2, Hb3, Hb4*** | Chr15 | 14187451 | 14186569 | - | | can be b2,3 or 4 |
| ***Hba2*** | Chr15 | 14185002 | 14184345 | + | | |
| ***Hb2, Hb3, Hb4*** | Chr15 | 14182238 | 14181356 | - | | can be b2,3 or 4 |
| ***Hba2*** | Chr15 | 14179755 | 14179098 | - | | |
| ***Hba3*** | Chr15 | 14173445 | 14171291 | - | | |
| ***Hba2*** | Chr15 | 14169193 | 14169850 | + | | |
| ***Hb3, Hb4*** | Chr15 | 14166907 | 14167581 | + | | |
| ***LCMT1*** | Chr15 | 14148838 | 14153907 | + | 99-100 | |
| ***ARHGAP17*** | Chr15 | 14146940 | 14127350 | + | 99-100 | |

**Table S6.** Tests for inversion genotypes and Hardy Weinberg equilibrium, and estimates of $F_{IS}$ (calculated where we have deviation from HWE) within each of the delineated sub-clusters, for each listed inversion.

| Subcluster1 | Chi2 | Df | PrChi2 | PrExact | $F_{IS}$ |
|---|---|---|---|---|---|
| Bchr10 | 0,03240741 | 1 | 0,85713642 | 1 | 0,028 |
| Bchr12.01 | 0,69771852 | 1 | 0,40355137 | 0,383 | 0,129 |
| Bchr13 | 0,27726563 | 1 | 0,59849905 | 0,647 | 0,081 |
| Bchr14 | 3,37166667 | 1 | 0,06632662 | 0,129 | -0,283 |
| Bchr16.01 | 3,37166667 | 1 | 0,06632662 | 0,124 | -0,283 |
| Bchr17 | 0,14388447 | 1 | 0,70444946 | 0,537 | 0,059 |
| Bchr3 | 0,105 | 1 | 0,74591 | 1 | -0,050 |
| Bchr4.01 | 2,06373879 | 1 | 0,15083968 | 0,256 | 0,222 |
| Bchr4.02 | 5,09069959 | 1 | 0,02405449 | 0,047 | -0,348 |
| Bchr7 | 1,16666667 | 1 | 0,28008721 | 0,596 | -0,167 |
| Bchr9 | 0,46537396 | 1 | 0,4951231 | 1 | -0,105 |
| Bchr2 | 3,4460964 | 1 | 0,06340134 | 0,12 | -0,286 |
| Bchr12.02 | 1,21909465 | 1 | 0,2695384 | 0,311 | 0,170 |
| Bchr18 | 0,47399371 | 1 | 0,49115559 | 1 | -0,106 |
| **Subcluster2** | **Chi2** | **Df** | **PrChi2** | **PrExact** | $F_{IS}$ |
| Bchr10 | 1,64642297 | 1 | 0,19944648 | 0,234 | 0,106 |
| Bchr12.01 | 2,50368481 | 1 | 0,11358027 | 0,12 | 0,131 |
| Bchr13 | 4,79259638 | 1 | 0,02858231 | 0,045 | -0,181 |
| Bchr14 | 2,45200513 | 1 | 0,11737485 | 0,124 | -0,130 |
| Bchr16.01 | 3,11790532 | 1 | 0,07743627 | 0,11 | -0,146 |
| Bchr17 | 1,24694684 | 1 | 0,26413642 | 0,273 | 0,092 |
| Bchr3 | 6,73040914 | 1 | 0,00947829 | 0,014 | 0,215 |
| Bchr4.01 | 0,30257939 | 1 | 0,58226987 | 0,689 | -0,046 |
| Bchr4.02 | 1,94279715 | 1 | 0,16336515 | 0,211 | -0,115 |

| | | | | | |
|---|---|---|---|---|---|
| Bchr7 | 0,15106705 | 1 | 0,69751773 | 1 | -0,032 |
| Bchr9 | 0,15298603 | 1 | 0,69569798 | 1 | -0,033 |
| Bchr2 | 0,00042662 | 1 | 0,98352104 | 1 | 0,002 |
| Bchr12.02 | 2,30258949 | 1 | 0,12915851 | 0,147 | -0,126 |
| Bchr18 | 0,01775957 | 1 | 0,89398378 | 0,766 | 0,011 |
| **Cluster3** | **Chi2** | **Df** | **PrChi2** | **PrExact** | **FIS** |
| Bchr10 | 3,01451839 | 1 | 0,08252196 | 0,091 | 0,217 |
| Bchr12.01 | 1,30612245 | 1 | 0,25309791 | 0,247 | 0,143 |
| Bchr13 | 0,40525573 | 1 | 0,52438738 | 0,736 | -0,080 |
| Bchr14 | 1,26913679 | 1 | 0,25992868 | 0,287 | -0,141 |
| Bchr16.01 | 3,11111111 | 1 | 0,0777599 | 0,116 | 0,222 |
| Bchr17 | 0,74895156 | 1 | 0,38680838 | 0,456 | 0,108 |
| Bchr3 | 1,30112772 | 1 | 0,25400734 | 0,322 | 0,143 |
| Bchr4.01 | 0,11880348 | 1 | 0,73033585 | 0,775 | 0,043 |
| Bchr4.02 | 0,65564097 | 1 | 0,41810308 | 0,607 | -0,101 |
| Bchr7 | 0,96522007 | 1 | 0,32587518 | 1 | -0,123 |
| Bchr9 | 1,10358997 | 1 | 0,2934796 | 0,464 | -0,132 |
| Bchr2 | 0,0542581 | 1 | 0,8158129 | 0,805 | 0,029 |
| Bchr12.02 | 0,01827273 | 1 | 0,89247221 | 1 | 0,017 |
| Bchr18 | 1,05193951 | 1 | 0,30506082 | 0,433 | -0,128 |
| **Subcluster4** | **Chi2** | **Df** | **PrChi2** | **PrExact** | |
| Bchr10 | 0,02267574 | 1 | 0,88030337 | 1 | |
| Bchr12.01 | 0,02770083 | 1 | 0,86781411 | 1 | |
| Bchr13 | 1,11111111 | 1 | 0,29184055 | 1 | |
| Bchr14 | 0,02267574 | 1 | 0,88030337 | 1 | |
| Bchr16.01 | 0,02267574 | 1 | 0,88030337 | 1 | |
| Bchr17 | 0,09781427 | 1 | 0,75446853 | 1 | |
| Bchr3 | 0,4 | 1 | 0,52708926 | 0,565 | |
| Bchr4.01 | 0,0010203 | 1 | 0,97451815 | 1 | |
| Bchr4.02 | 0,625 | 1 | 0,4291953 | 1 | |
| Bchr7 | 0,12345679 | 1 | 0,72531515 | 1 | |
| Bchr9 | 1,11111111 | 1 | 0,29184055 | 1 | |
| Bchr2 | 0,4 | 1 | 0,52708926 | 1 | |
| Bchr12.02 | 2,74376417 | 1 | 0,09763454 | 0,132 | |
| Bchr18 | 1,11111111 | 1 | 0,29184055 | 1 | |
| **Subcluster5** | **Chi2** | **Df** | **PrChi2** | **PrExact** | |
| Bchr10 | 2,99022222 | 1 | 0,08376867 | 0,169 | |
| Bchr12.01 | 0,01314924 | 1 | 0,90870657 | 1 | |
| Bchr13 | 0,078125 | 1 | 0,77985462 | 1 | |
| Bchr14 | 0,04535147 | 1 | 0,83135906 | 1 | |
| Bchr16.01 | 0,8 | 1 | 0,37109337 | 0,565 | |
| Bchr17 | 0,89990817 | 1 | 0,34280633 | 1 | |

| | | | | |
|---|---|---|---|---|
| Bchr3 | 0,89990817 | 1 | 0,34280633 | 0,665 |
| Bchr4.01 | 1,28586619 | 1 | 0,25681116 | 0,618 |
| Bchr4.02 | 0,032 | 1 | 0,85802766 | 1 |
| Bchr7 | 1,9755102 | 1 | 0,1598642 | 0,235 |
| Bchr9 | 2,87752675 | 1 | 0,08982389 | 0,272 |
| Bchr2 | 0,21117958 | 1 | 0,64584443 | 1 |
| Bchr12.02 | 0,03472222 | 1 | 0,85217893 | 1 |
| Bchr18 | 0,01314924 | 1 | 0,90870657 | 1 |

**Table S7.** Sexing of polar cod by genotyping. Specimen ID, sex by morphology (visual inspection of gonads), single nucleotide polymorphisms (SNPs) within genes displaying male/female linked variants, and the final decided sex for each sample given. In the table "Female" denotes 0/0, and "Male" 0/1 variants.
Table provided online, as an Excel sheet:
https://www.mn.uio.no/ibv/personer/vit/snhoff/table_s7_sex_by-genotyping.xlsx

**Table S8.** Summary of number of sequenced specimens from each sample location.

| **Sample location** | **Samples (n)** |
|---|---|
| Northeast Barents Sea autumn 2019 | 58 |
| Northwest Barents Sea summer 2018 (JC1) | 60 |
| Northwest Barents Sea summer 2019 (Q3) | 32 |
| Northwest Barents Sea winter 2019 (Q4) | 26 |
| Billefjorden | 22 |
| Hornsund | 20 |
| Iceland | 24 |
| Kongsfjorden | 20 |
| 0-group | 20 |
| Davis strait, Canada | 4 |
| Tyrolerfjorden, Greenland | 4 |

**Tables S9-17.** Information about specimens sequenced including sample location, date of sampling, and recorded metadata.
Table provided online, as an Excel sheet:
https://www.mn.uio.no/ibv/personer/vit/snhoff/table_s9-s17_metadata_290_samples.xlsx

**Table S18.** Overview of numbers of SNPs called and retained after filtering among 290 polar cod.

| Chromosome | Raw SNPs | SNPs that "PASS" hard filtering | Final quality filtered SNPs: QD5, biallelic, MAF 0.01 |
|---|---|---|---|
| Chr1 | 2 208 739 | 1 520 200 | 231 714 |
| Chr2 | 2 069 968 | 1 402 780 | 229 738 |
| Chr3 | 2 085 521 | 1 378 726 | 222 421 |
| Chr4 | 1 927 828 | 1 272 072 | 192 197 |
| Chr5 | 1 630 177 | 1 125 778 | 166 025 |
| Chr6 | 1 942 283 | 1 173 874 | 155 501 |
| Chr7 | 1 281 131 | 905 845 | 131 181 |
| Chr8 | 1 269 229 | 919 509 | 141 205 |
| Chr9 | 1 377 761 | 1 036 057 | 158 413 |
| Chr10 | 1 342 616 | 968 081 | 149 134 |
| Chr11 | 1 137 902 | 827 989 | 115 641 |
| Chr12 | 1 082 856 | 746 345 | 115 148 |
| Chr13 | 1 113 077 | 823 731 | 132 531 |
| Chr14 | 1 141 791 | 821 992 | 119 634 |
| Chr15 | 1 009 409 | 753 336 | 122 199 |
| Chr16 | 1 004 111 | 699 419 | 96 509 |
| Chr17 | 1 001 320 | 726 054 | 112 415 |
| Chr18 | 1 008 083 | 730 560 | 113 419 |
| Mt-contig | 1309 | | |
| Combined total | 25 635 111 | 17 832 348 | 2 705 025 |

**Table S19.** Genomic intervals that were removed to create a putative neutral dataset with regard to linked chromosomal regions.

| Chromosome | Start region removed (bp) | End region removed (bp) | Note |
|---|---|---|---|
| Chr1 | 14500000 | 18000000 | |
| Chr1 | 22000000 | 36000000 | |
| Chr10 | 1 | 9500000 | |
| Chr10 | 14500000 | 20500000 | |
| Chr11 | 5500000 | 18500000 | |
| Chr12 | -499999 | 4000000 | remove whole chromosome |
| Chr12 | 12000000 | 15100000 | |
| Chr13 | 1 | 4500000 | |
| Chr13 | 1 | 16500000 | more than half chromosome |
| Chr14 | 4500000 | 11500000 | |
| Chr14 | 500000 | 5500000 | |
| Chr15 | 8500000 | 15500000 | more than half 9mb-end |
| Chr16 | 1900000 | 3200000 | |
| Chr16 | 4500000 | 10500000 | |
| Chr17 | 6000000 | 12500000 | more than half chr 6mb-end |
| Chr18 | 1 | 5500000 | |
| Chr18 | 8000000 | 10000000 | |
| Chr2 | 22500000 | 40500000 | |
| Chr3 | 33700000 | 36500000 | |
| Chr3 | 1 | 18500000 | |
| Chr4 | 1340000 | 2650000 | |
| Chr4 | 2500000 | 4500000 | |
| Chr4 | 17500000 | 27500000 | |
| Chr5 | 9500000 | 30500000 | |
| Chr6 | 13500000 | 26000000 | |
| Chr7 | 1 | 3500000 | |
| Chr7 | 9500000 | 20500000 | |
| Chr8 | 10500000 | 220500000 | |
| Chr9 | 21550000 | 241500000 | |
| Chr9 | 5500000 | 15500000 | |
